# Developmentally-regulated impairment of parvalbumin interneuron synaptic transmission in an experimental model of Dravet syndrome

**DOI:** 10.1101/2021.07.28.454042

**Authors:** Keisuke Kaneko, Christopher B. Currin, Kevin M. Goff, Ala Somarowthu, Tim P. Vogels, Ethan M. Goldberg

**Author notes:** Correspondence: Ethan M. Goldberg, M.D., Ph.D.,The Children’s Hospital of Philadelphia, Abramson Research Center, Room 502A, Philadelphia, PA 19104.

## Abstract

Dravet syndrome (DS) is a neurodevelopmental disorder defined by epilepsy, intellectual disability, and sudden death, due to heterozygous variants in SCN1A with loss of function of the sodium channel subunit Nav1.1. Nav1.1-expressing parvalbumin GABAergic interneurons (PV-INs) from pre-weanling Scn1a+/− mice show impaired action potential generation. A novel approach assessing PV-IN function in the same mice at two developmental time points showed that, at post-natal day (P) 16-21, spike generation was impaired all mice, deceased prior or surviving to P35. However, synaptic transmission was selectively dysfunctional in pre-weanling mice that did not survive. Spike generation in surviving mice normalized by P35, yet we again identified abnormalities in synaptic transmission. We conclude that combined dysfunction of PV-IN spike generation and synaptic transmission drives disease severity, while ongoing dysfunction of synaptic transmission contributes to chronic pathology. Modeling revealed that PV-IN axonal propagation is more sensitive to decreases in sodium conductance than spike generation.

## Introduction

Dravet syndrome (DS) is a neurodevelopmental disorder defined by temperature-sensitive seizures, treatment-resistant epilepsy, developmental delay/intellectual disability with features of autism spectrum disorder (ASD), and increased incidence of sudden unexpected death (SUDEP). DS is due to heterozygous pathogenic loss-of-function variants in the gene SCN1A encoding the voltage-gated sodium (Na+) channel *α* subunit Nav1.1 [Dravet, 2011, Scheffer, 2012]. While new pharmacological therapies for DS have emerged including cannabidiol [Kaplan et al., 2017, Khan et al., 2018] and fenfluramine [Ceulemans et al., 2016, Schoonjans et al., 2017] and antisense oligonucleotide-based approaches may be on the horizon [Han et al., 2020], treatment remains essentially palliative, and there is no cure. Previous studies in experimental mouse models of DS (Scn1a+/− mice) showed prominent dysfunction of GABAergic interneurons (INs) in the neocortex and hippocampus at early developmental time points (second and third postnatal weeks), particularly the parvalbumin (PV)-expressing subclass (PV-INs) [Yu et al., 2006, Ogiwara et al., 2007, Tai et al., 2014, De Stasi et al., 2016]. However, recent data suggests that impairments of spike generation may be transient and limited to early development [Favero et al., 2018], which raises the question as to the mechanism of continued epilepsy, intellectual disability, ASD, and sudden death seen in Scn1a+/− mice and DS patients. This finding also has important implications for emerging efforts to increase expression of SCN1A as a potential therapeutic strategy [Colasante et al., 2019, Han et al., 2020, Yamagata et al., 2020]; upregulation of SCN1A may be ineffective or even deleterious if administered after disease onset if chronic pathology is driven by an alternative mechanism after initial compensatory molecular or circuit rearrangements.

Accurate assessment of PV-IN function across development is confounded by the fact that most data on PV-IN function/dysfunction in DS derives from recordings from acutely dissociated neurons or brain slices from Scn1a+/− mice before or at onset of epilepsy (prior to P21), yet a large proportion of these mice subsequently are deceased from status epilepticus or a SUDEP-like phenomenon. It is possible – and perhaps likely – that mice remaining alive represent a nonrandom subset of the initial population. It is well known that there is a spectrum of disease severity in Scn1a+/− mice [Hawkins et al., 2016, 2017], similar to that observed in humans [Harkin et al., 2007, Mei et al., 2019], the basis of which is unclear but may involve genetic modifiers [Hawkins and Kearney, 2016, Hawkins et al., 2016, Calhoun et al., 2017]. This concern may pertain to any study involving an experimental animal model of a neurodevelopmental disorder associated with mortality. Restated: if we could retrospectively record PV-INs at early time points (P16-21) in those Scn1a+/− mice where PV-IN function was later determined to be normal (e.g., P35-56), would these cells be normal or abnormal? Conversely, would PV-INs in those Scn1a+/− mice deceased at early time points (< P35) have exhibited functional recovery if these mice had survived to and been assessed at/beyond P35? Here, we address this critical question using a novel experimental paradigm to assess PV-IN function in the same mice at both time points (< P21 and > P35) in “mini-slices” from young (P16-21) Scn1a.PV-Cre.tdTomato mice, and then again in acute brain slices at later time points (P35-56) from those mice that remained alive, recording a comprehensive panel of cellular and synaptic properties. We found impaired action potential generation at early time points (P16-21) in PV-INs from Scn1a+/− mice that did and did not survive until at least P35 (i.e., all mice).

We also reconcile the apparent normalization of PV-IN spike generation with the ongoing epilepsy, behavioral and cognitive impairment, and continued mortality observed in Scn1a+/− mice during the chronic phase of the disorder. We examined unitary synaptic transmission at PV-IN synapses in Scn1a+/− vs. control mice across development by performing multiple whole-cell patch-clamp recordings of genetically identified PV-INs and target principal cells (PCs). We found severe impairments in synaptic transmission at PV-IN:PC synapses selectively in those P16-21 Scn1a+/− mice that were subsequently deceased prior to P35, but not in those surviving to P35-56. Spike generation of PV-INs from Scn1a+/− mice surviving beyond P35 was found to have normalized by P35-56, as described previously [Favero et al., 2018]; yet, we again identified dysfunction of synaptic transmission at PV-IN:PC synapses at this later time point. Photoactivation of a somatically-targeted excitatory opsin expressed in PV-INs reproduced the genotype-specific abnormality in neurotransmission while activation of a synaptically-localized opsin bypassed the presumptive axonal dysfunction. Results of a multicompartmental model of a PV-IN supported the conclusion that action potential propagation is more sensitive to decreased Na+ conductance than spike generation, and highlighted potential mechanisms for the selective recovery of spike generation in PV-INs in young adult Scn1a+/− mice.

These findings are consistent with PV-IN axonal dysfunction as a core pathomechanism of ongoing chronic DS and suggest that the safety factor for axonal conduction is less than for spike generation at the axon initial segment. Results highlight a potential marker of SUDEP risk and illuminate differential mechanisms of epilepsy onset versus chronic pathology in DS, shifting the locus of dysfunction from that of PV-IN spike generation to that of synaptic transmission.

## RESULTS

### Targeting PV Interneurons in Wild-type vs. Scn1a+/− Mice Across Development

We used a well-established mouse model of Dravet syndrome (Scn1a+/− mice) in which a loxP-flanked neo cassette replaces exon 1 of the Scn1a gene, creating a null allele [Mistry et al., 2014]. Mice were maintained on a strict 50:50 C57BL6/J:129S6 genetic background (see METHODS for details) which recapitulates salient core features of DS observed in human patients, including temperature-sensitive seizures, early-onset chronic epilepsy (with appearance of seizures at/around post-natal day (P) 18), approximately 30% death from seizure-related causes during early development (compared to 20% lifetime risk in humans) [Cooper et al., 2016], and social recognition and memory deficits consistent with features of ASD [Berkvens et al., 2015, He et al., 2018, Xiong et al., 2016]. Cross to double transgenic PV-Cre.tdTomato mice yielded triple transgenic Scn1a.PV-Cre.tdT mice and age-matched wild-type PV-Cre.tdT littermate controls [Favero et al., 2018] and facilitated comparison of PV-INs between genotypes and across development.

To facilitate recordings from PV-INs form the same mice at two developmental time points (pre-weanling and young adult), we developed a novel technique referred to here as “mini-slices” (see METHODS) generated from microbiopsies of primary somatosensory neocortex of young (P16-21) Scn1a.PV-Cre.tdTomato mice, followed by acute brain slice preparation using standard techniques at later time points (P35-56) from those mice that survived to young adulthood. Scn1a+/− mice were retrospectively defined as Group A if deceased prior to P35 and Group B if the animal survived to P35-56. Scn1a+/− mice surviving to and recorded at P35-56 were defined as the Group C dataset (Figure 1). Rate of spontaneous death was 26.2% (22 of 84) in Scn1a+/− mice (Group A, n = 22; Group B, n = 62), similar to the 25.1% rate in our previous study [Favero et al., 2018] and in the published literature (e.g., [Hawkins et al., 2017, Han et al., 2020]). The same mini-slice procedure in wild-type mice (pre-weanling, Group D; young adult, Group E) was associated with 0% mortality (0 of 37), indicating that this procedure was well-tolerated.

**Figure 1:**
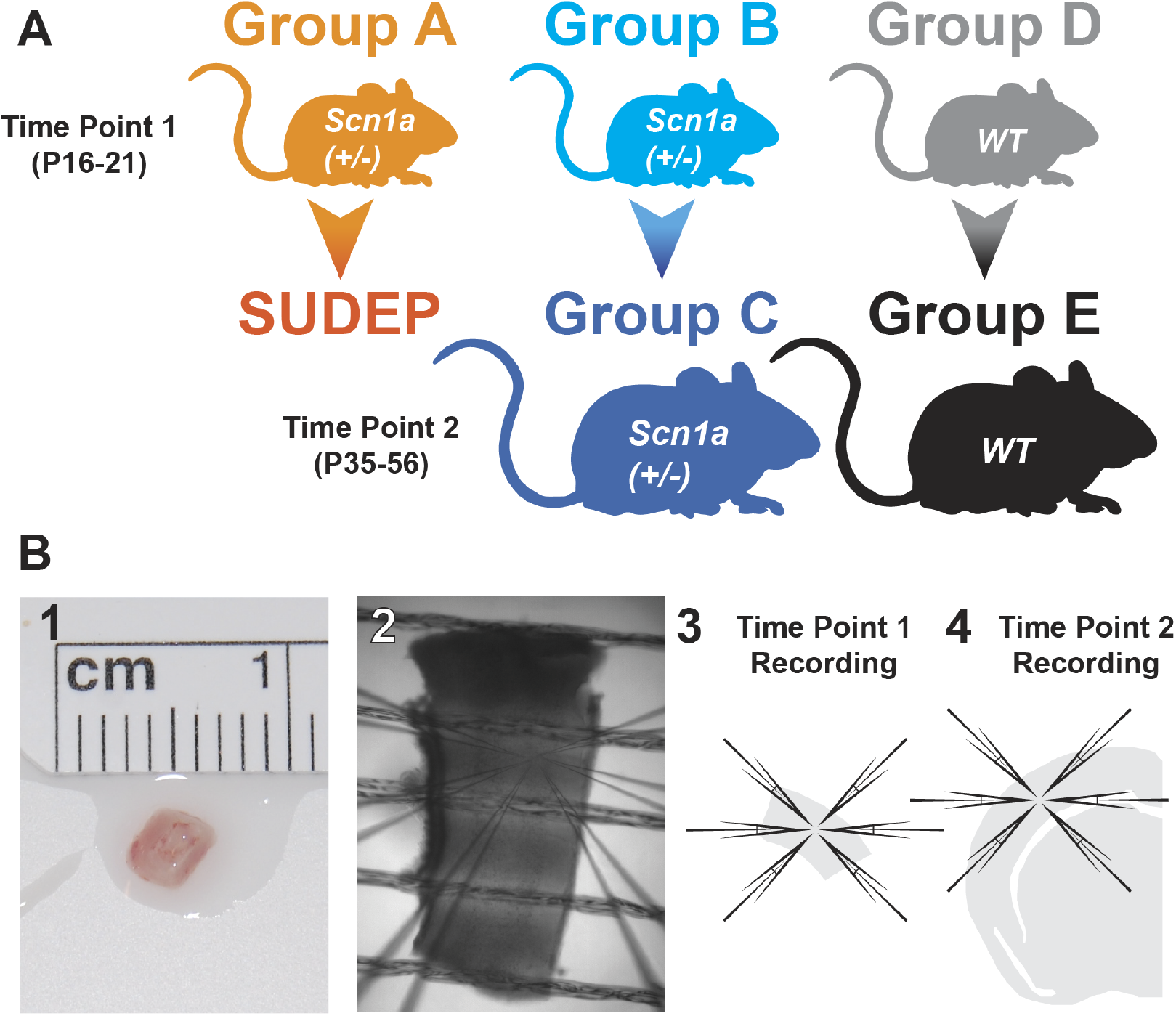
Analysis of PV interneurons across development. (A) Five experimental groups were studied: Scn1a+/− mice recorded at P16-21 and deceased by P35 (Group A; orange); Scn1a+/− mice recorded at P16-21 that survive to P35-56 (Group B; light blue); Scn1a+/− mice recorded at P35-56 (Group C; dark blue); wild-type littermates at P16-21 (Group D; gray) and P35-56 (Group E; black). (B) The mini-slice procedure. Mini-slices (B1-3) were generated from biopsies of primary somatosensory neocortex of pre-weanling (P16-21) Scn1a.PV-Cre.tdTomato and age-matched WT.PV-Cre.tdT mice. Acute brain slices (B4) were generated using standard techniques at later time points (P35-56) from Scn1a+/− mice that did not die prior to P35 (Group C) and from age-matched wild type littermates at P35-56 (Group E).

We then defined five experimental datasets: pre-weanling Scn1a+/− mice recorded at P16-21 that were deceased by P35 (Group A); pre-weanling Scn1a+/− mice recorded at P16-21 that survive to P35-56 (Group B), and young adult Scn1a+/− mice recorded at P35-56 (Group C; i.e., the same mice as in Group B, but recorded again at P35-56 to generate a separate dataset), as well as pre-weanling wild-type (WT) littermates at P16-21 (Group D) and P35-56 (Group E; the same mice as in Group D). We then focused on six key comparisons: Group A:Group B (to assess any difference between pre-weanling Scn1a+/− mice recorded at P16-21 that die prior to P35 vs. those that survive to young adulthood), A:D and B:D (to determine the difference between the two groups of pre-weanling Scn1a+/− mice vs. age-matched WT littermate controls), B:C (to assess the developmental trajectory of PV-IN excitability in surviving Scn1a+/− mice), C:E (to compare surviving young adult Scn1a+/− mice to age-matched WT controls), and D:E (to assess the developmental trajectory of PV-IN excitability in WT mice).

### PV-INs from Scn1a+/− Mice Exhibit Severe Impairments in Action Potential Generation at Early Time Points

We then performed whole-cell patch-clamp recordings from identified tdTomato-positive PV-INs in layer 4 primary somatosensory neocortex and measured a range of passive membrane properties, features of single action potentials, and properties of repetitive spiking.

We found that PV-INs recorded in pre-weanling Scn1a+/− mice at P16-21 that either were deceased by P35 (Group A; Figure 2A1) and PV-INs recorded at P16-21 from Scn1a+/− mice that survived to P35-42 (Group B; Figure 2A2) all exhibited markedly abnormal electrical excitability relative to age-matched wild-type littermate controls (Group D; wild-type mice at P16-21; Figure 2A4) and to PV-INs from young adult Scn1a+/− mice recorded at P35-56 (Group C; Figure 2A3), across a range of measures that depend on Na+ channel function, including indicators of action potential generation and repetitive firing. For example, maximal instantaneous firing frequency was 231.6 ± 14.2 Hz for pre-weanling Scn1a+/− mice P16-21 deceased prior to P35 (Group A; n = 21 cells from 6 mice) and 246.6 ± 10.7 Hz for pre-weanling Scn1a+/− mice that survived to P35 (Group B; n = 44 cells from 19 mice; ANOVA F(4,126) = 3.296; p = 0.364 vs. Group A; post-hoc correction for multiple comparisons with Holm-Sidak test), but was 324.8 ± 8.9 Hz for young adult Scn1a+/− mice at P35-56 (Group C; n = 35 cells from 12 mice; p < 0.0001 vs. Group B), 287.4 ± 6.6 Hz for pre-weanling WT mice (Group D; n = 28 cells from 9 mice; p = 0.006 vs. Group A; p = 0.014 vs. Group B) and 340.8 ± 11.3 Hz for young adult WT mice (Group E; n = 17 cells from 9 mice; p = 0.57 vs. Group C; p = 0.01 vs. Group D). PV-INs from WT mice exhibited higher firing frequency at the later time point (Group E relative to Group D), consistent with the normal developmental maturation of PV-IN excitability [Goldberg et al., 2011]. However, PV-INs in all pre-weanling Scn1a+/− mice (Group A and B), irrespective of the subsequent occurrence of SUDEP, exhibited markedly decreased excitability relative to both PV-INs from age-matched WT controls (Group D) as well as PV-INs from Scn1a+/− mice that survived to young adulthood (Group C) (Figure 2B-C; Table S2). Identical results were obtained when measuring firing frequency at two times rheobase current injection (to normalize for differences in input resistance; Table S2; see METHODS), and the same between group differences were also observed for other electrophysiological properties known to depend on Na+ current density, including rate of rise of the action potential (dV/dt), action potential half-width (AP ½-width), and other measures that do not directly depend on Na+ channels (see Table S3). These genotype- and age-dependent group differences in firing frequency were observed for both male and female mice (not shown). To confirm the basic finding that action potential generation by PV-INs in pre-weanling mice surviving to young adulthood (Group B) was in fact the same as for those in mice decreased by P35 (Group A) yet different than age-matched wild-type controls (Group D), we constructed a linear mixed model to account for variability between mice. The mixed model was fit to the data with mouse as a random factor, which confirmed that Group B was significantly different than Group D but the same as Group A.

**Figure 2:**
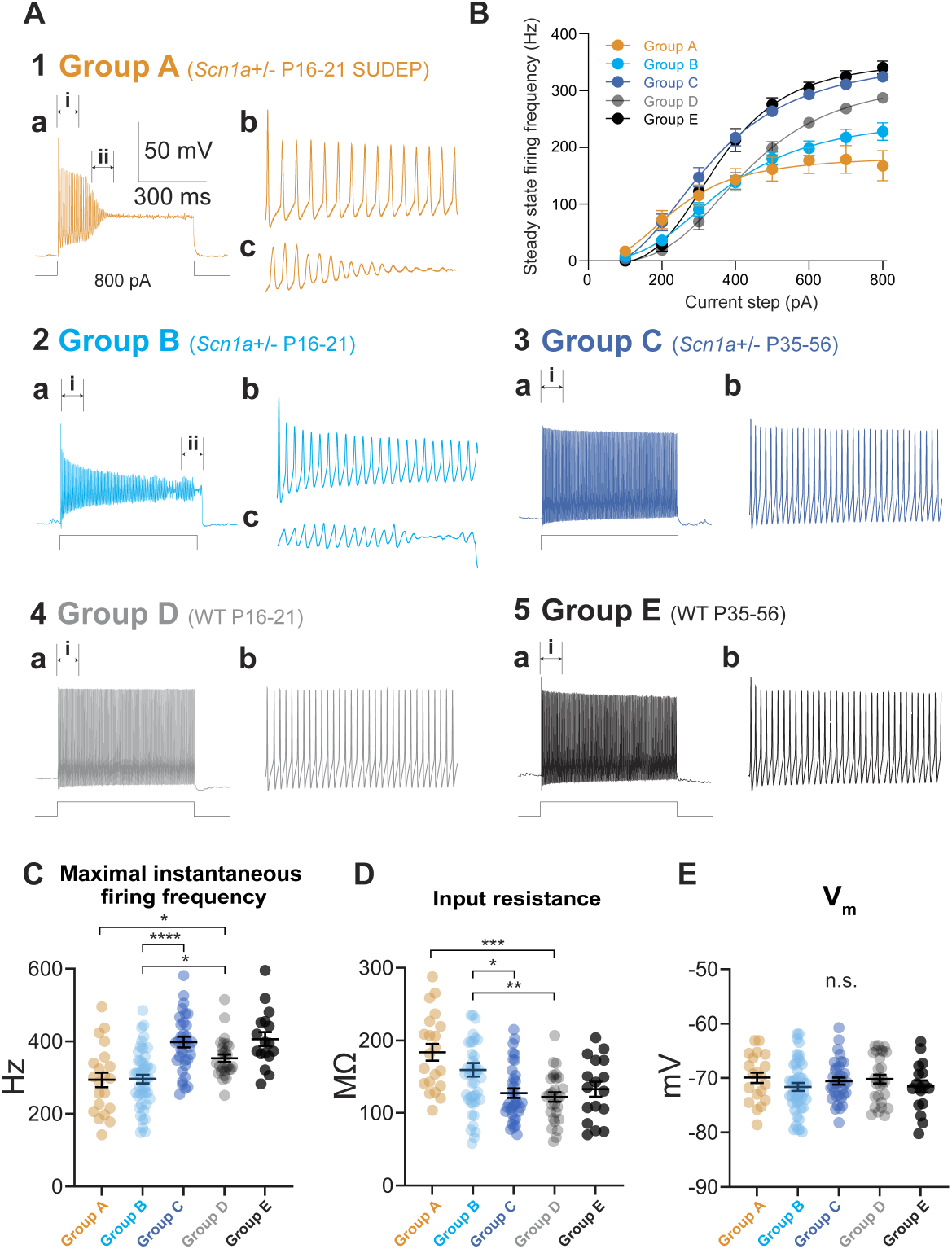
Transient impairments in action potential generation in PV interneurons in all Scn1a+/− mice. (A) Representative current clamp recordings from PV-INs from all experimental Groups A-E (A1-5). Note that Groups A and A exhibited markedly abnormal excitability with action potential failure (shown in detail in A1a-c and A2a-c), while Group C exhibits normal action potential generation and repetitive firing similar to Group E. (B) Input–output curves showing steady state firing frequency for PV-INs from each group. Excitability of pre-weanling Scn1a+/− mice that did not (Group A) and did survive to young adulthood (Group B) was decreased relative to age-matched WT (Group D; *; p < 0.05) and to PV-INs from young adult Scn1a+/− mice (Group C; ****, p < 0.0001). Curves for the WT pre-weanling mice at P16-21 was also different from young adult WT (Group E; ****; p < 0.0001). See Figure S2 for separate plots depicting individual comparisons. (C-E) Summary data for maximal instantaneous firing frequency (C), input resistance (D), and resting membrane potential (E) in PV-INs from each group. , p < 0.05; **, p < 0.01; ***, p < 0.001; ****, p < 0.0001 via one-way ANOVA with post-hoc correction for multiple comparisons via Holm-Sidak test.

In contrast, we found no differences in passive membrane properties such as resting membrane potential and membrane potential sag (Figure 2E; Figure S3C; Table S1), although we did find genotype differences in input resistance (Figure 2D; Table S1) and measures related to this (rheobase current injection and membrane time constant; Figure S3A and Figure S3B; Table S1).

This data demonstrates that action potential generation is impaired at early time points in all PV-INs from Scn1a+/− mice at < P21, including those that do and do not survive to P35, while confirming our prior results [Favero et al., 2018] that action potential generation of PV-INs from Scn1a+/− mice has normalized by P35-56.

### Group-specific Impairments in PV Interneuron Synaptic Transmission in Scn1a+/− Mice

The findings reported thus far represent somewhat of a paradox and raise two critical questions relevant to pathomechanisms of DS. Action potential generation and repetitive/high-frequency firing of PV-INs was impaired in both Scn1a+/− Group A and B, yet Group A mice were deceased by P35 while Group B mice survived to P35-56. The basis for this profound difference is unclear. Second, PV-INs from young adult Scn1a+/− mice Group C were found to have recovered high frequency discharge capabilities characteristic of mature fast-spiking PV-INs [Jonas et al., 2004, Hu et al., 2014, Favero et al., 2018] and were near-equivalent to age-matched WT controls (Group E). However, Scn1a+/− mice that survive to P35 continue to exhibit chronic epilepsy, cognitive impairment, features of ASD, and ongoing SUDEP. This indicates that the basis of chronic pathology in DS cannot be abnormal PV-IN spike generation. We hypothesized that, while PV-INs may have recovered normal or near-normal capacity for spike generation, persistent deficits might exist in synaptic transmission – perhaps due to impaired spike propagation –given the known dependence of PV-IN axonal function on a high Na+ channel density at the AIS [Hu and Jonas, 2014].

To assay the fidelity of GABAergic transmission at PV-IN synapses, we performed multiple whole-cell patch-clamp recordings from identified tdTomato-positive PV-INs as well as principal cells in layer 4 primary somatosensory neocortex and measured a range of properties of evoked inhibitory post-synaptic currents recorded in PCs (IPSCs; in voltage clamp) in response to action potentials in presynaptic PV-INs (generated in current clamp; Figure 3A). PCs were patched using an internal solution containing high (63 mM) chloride (Cl) such that E_Cl_*−* = -17 mV and IPSPs were recorded as depolarizations from resting membrane potential, while IPSCs were recorded as large inward currents at a holding potential of -80 mV (see METHODS). This approach facilitated detection of small IPSCs versus synaptic failures, which was critical for subsequent analysis.

**Figure 3:**
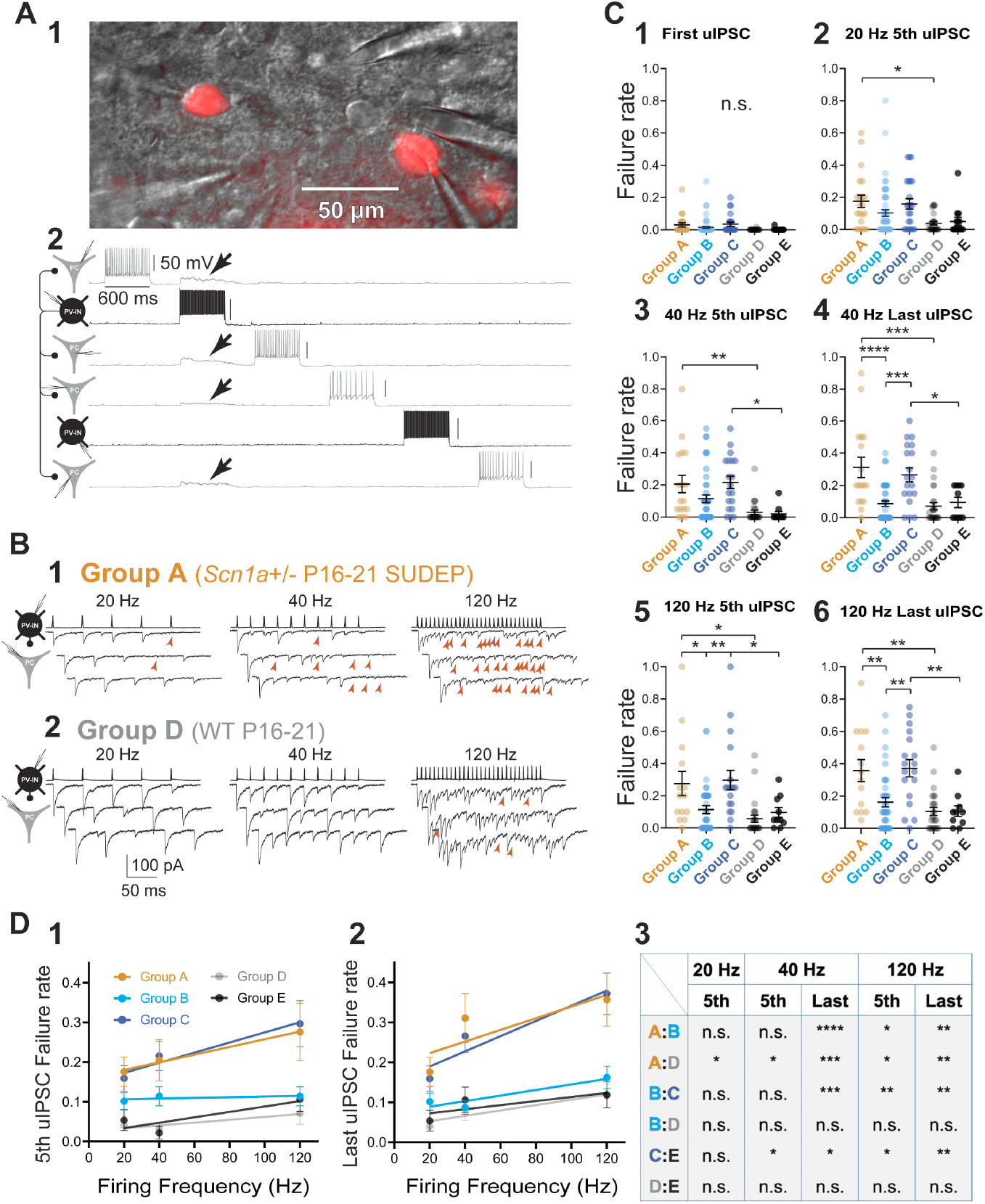
Persistent impairments in AP propagation in PV interneurons. (A) Multiple simultaneous whole-cell recording from identified tdTomato-positive PV-INs as well as principal cells in layer 4 primary somatosensory cortex. The example shown is from a pre-weanling WT mouse (Group D). Shown is a representative image of the multiple whole-cell recording of two PV-INs (cells #2 and #5) and four PCs (cells #1, #3, #4, #6) in the mini-slice preparation under infrared microscopy differential interference contrast and corresponding epifluorescence image (A1; showing tdTomato-positive PV-INs). Four synaptic connections were identified from Cell #2 (in black) which produce large depolarization in Cells #1, #2, #3, and #4 (arrows; in gray; A2). (B) Representative examples of evoked unitary inhibitory postsynaptic currents (uIPSCs) at 20, 40, 120 Hz in Group A (B1) and Group D (B2). Presynaptic PV-INs were recorded in current-clamp and postsynaptic PCs were recorded in voltage-clamp. IPSCs were recorded as large inward currents. Red arrowheads indicate synaptic failures. (C) Summary data for the failure rate of the 1st (C1) and last (i.e., 5th; C2) uIPSCs at 20 Hz, 5th (C3) and last (i.e., 10th; C4) uIPSCs at 40 Hz, and 5th (C5) and last (i.e., 30th; C6) uIPSCs at 120 Hz. Failure rate (range, 0 to 1) is calculated as the number of failures divided by total stimuli (failures plus number of success). See Table S4. Note that failures occurred at frequencies that PV-INs could clearly generate at the soma and was more pronounced with successive stimulation (Figure 3C4, 6) yet was still present early in trains (Figure 3C3, 5). (D) Linear fit of the relationship between failure rate and presynaptic firing frequency for the 5th (D1) and last uIPSC (D2) in trains. See Table S4. This relationship for pre-weanling Scn1a+/− mice that did not survive to young adulthood (Group A) was significantly different vs. pre-weanling Scn1a+/− mice that did survive (Group B; ***, p < 0.001 vs. Group A) and age-matched WT mice (Group D; ****, p < 0.0001 vs. Group A). Data from surviving young adult Scn1a+/− mice (Group C) was significantly different from age-matched WT (Group E; ****; p < 0.0001) (D3).

The PV-IN:PC synapse is known to be a high-fidelity connection with a low (or near-zero) failure rate due to high release probability from multiple boutons per synaptic connection [Galarreta and Hestrin, 1998, Kraushaar and Jonas, 2000, Ali et al., 2001, Beierlein et al., 2003, Goldberg et al., 2005, Hu et al., 2014]. We stimulated presynaptic PV-INs with single action potentials and found a very low rate of synaptic failures at PV-IN:PC connections in WT mice in both pre-weanling (Group D; 0.001 ± 0.001; n = 15 connections from 7 mice) and young adult WT mice (Group E; 0.002 ± 0.002; n = 11 connections from 9 mice), consistent with published results. However, we detected failures at PV-IN:PC connections in Scn1a+/− mice, including Group A (0.03 ± 0.01; n = 17 connections from 7 mice; p = 0.151 vs. Group D) and Group C (0.03 ± 0.01; n = 18 connections from 15 mice; p = 0.191 vs. Group E), with a lower but still non-zero failure rate in Group B (0.01 ± 0.01; n = 43 connections from 22 mice; p = 0.432 vs. Group A and 0.432 vs. Group D).

We then stimulated presynaptic PV-INs to discharge trains of repetitive action potentials at various frequencies (20, 40, 120 Hz) and identified a marked increase in frequency-/activity-dependent synaptic failure at PV-IN:PC synapses specifically in Scn1a+/− mice Groups A and C, but not B (Figure 3B-C; Table S4). For example, in approximately one-third of trials, the final stimulus in a 10-pulse train at 40 Hz resulted in failure at PV-IN:PC connections in pre-weanling Scn1a+/− mice that were deceased prior to P35 (Group A; 0.31 ± 0.06; n = 21 connections from 7 mice), a rate that was approximately three-fold higher than for synapses in developing Scn1a+/− mice at P16-21 that survive to P35-56 (Group B; 0.09 ± 0.02; n = 48 connections from 22 mice; ANOVA F(4, 103) = 2.957; p < 0.0001 vs. Group A with post-hoc correction for multiple comparisons with Holm-Sidak test) or for age-matched WT littermate controls (Group D; 0.08 ± 0.03; n = 21 connections from 9 mice; p = 0.0002 vs. Group A). Failure rate was also three-fold higher in young adult Scn1a+/− mice (Group C; 0.27 ± 0.04; n = 23 connections from 15 mice) than for age-matched WT littermate controls (Group E; 0.11 ± 0.03; n = 12 connections from 10 mice; p = 0.04 vs. Group C) (Figure 3Cd). For the last event in a 30-pulse train at 120 Hz, failures also occurred in approximately one-third of trials at PV-IN:PC synapses from surviving Scn1a+/− mice (Group C), which was 2.5-fold higher than for age-matched WT controls (Figure 3C6).

These synaptic failures may be due to failures of action potential propagation for the following reasons. First, the presence of a presynaptic action potential could be directly confirmed in the multiple whole-cell recording configuration, ruling out failure of spike generation (which was never apparent in PV-INs from Scn1a+/− mice except during repetitive high-frequency firing in response to prolonged, large-amplitude rectangular current injections). Second, the PV-IN:PC connection is characterized by large-amplitude IPSCs which were further augmented by our experimental conditions (i.e., high internal chloride concentration), and this – combined with < 2 pA peak-to-peak noise of our recording system – makes it unlikely that we were incorrectly labeling small IPSCs as failures. Finally, to further differentiate between failures of propagation vs. synaptic vesicle release, we performed an analysis of the contingency of the amplitude of the IPSC in response to the second presynaptic action potential (IPSC2) based on the success or failure of the first event (IPSC1) in a paired-pulse paradigm (Figure S6A). Action potential invasion of the presynaptic terminal leads to spike-evoked calcium influx which engages short-term synaptic plasticity mechanisms (in the case of the PV-IN synapse, this is synaptic depression; Zucker and Regehr, 2002; Xiang et al., 2002; Beierlein et al., 2003); hence, presence or absence of synaptic depression of IPSC2 can be used as an indicator of whether calcium influx occured and hence whether a somatic action potential arrived successfully at the presynapse. The amplitude of IPSC2 was smaller if IPSC1 was a success versus if IPSC1 was a failure (31.92 ± 7.03 pA for success vs. 48.75 ± 11.04 pA for failure; n = 20; p = 0.0023; Figure S6Ba), consistent with short-term synaptic depression (PPR = 0.65). However, if the response to the first presynaptic action potential resulted in failure, the amplitude of IPSC2 was the same as the mean amplitude of IPSC1 when IPSC1 was a success (48.75 ± 11.04 pA for IPSC2 if IPSC1 was a failure vs. 54.67 ± 10.76 pA for IPSC1 if IPSC1 was a success; n = 20; p = 0.17; Figure S6Bc). This data suggests that, in trials in which a failure occured, presynaptic calcium channels did not open, consistent with failure of propagation of the somatic action potential to the synapse.

In addition, we found that synaptic latency – the timing of synaptic transmission between connected neurons (measured from the peak of action potential to onset of the unitary IPSC; METHODS) – was prolonged in PV-INs from pre-weanling Scn1a+/− mice deceased by P35 (Group A) relative to age-matched WT littermate controls (Group D) and in PV-INs from young adult Scn1a+/− mice at P35-56 (Group C) vs. controls (Group E) (Figure 4A, Figure 4B, and Table S4), being 0.91 ± 0.06 ms in Group A (n = 21 connections from 7 mice), 0.81 ± 0.04 in Group B (n = 48 connections from 22 mice; ANOVA (4, 123) = 2.933; p = 0.25 vs. Group A), 0.86 ± 0.07 ms in Group C (n = 23 connections from 15 mice), 0.65 ± 0.03 in Group D (n = 21 connections from 9 mice; p = 0.004 vs. Group A; p = 0.06 vs. Group B), and 0.59 ± 0.03 ms in Group E (p = 0.01 vs. Group C).

**Figure 4:**
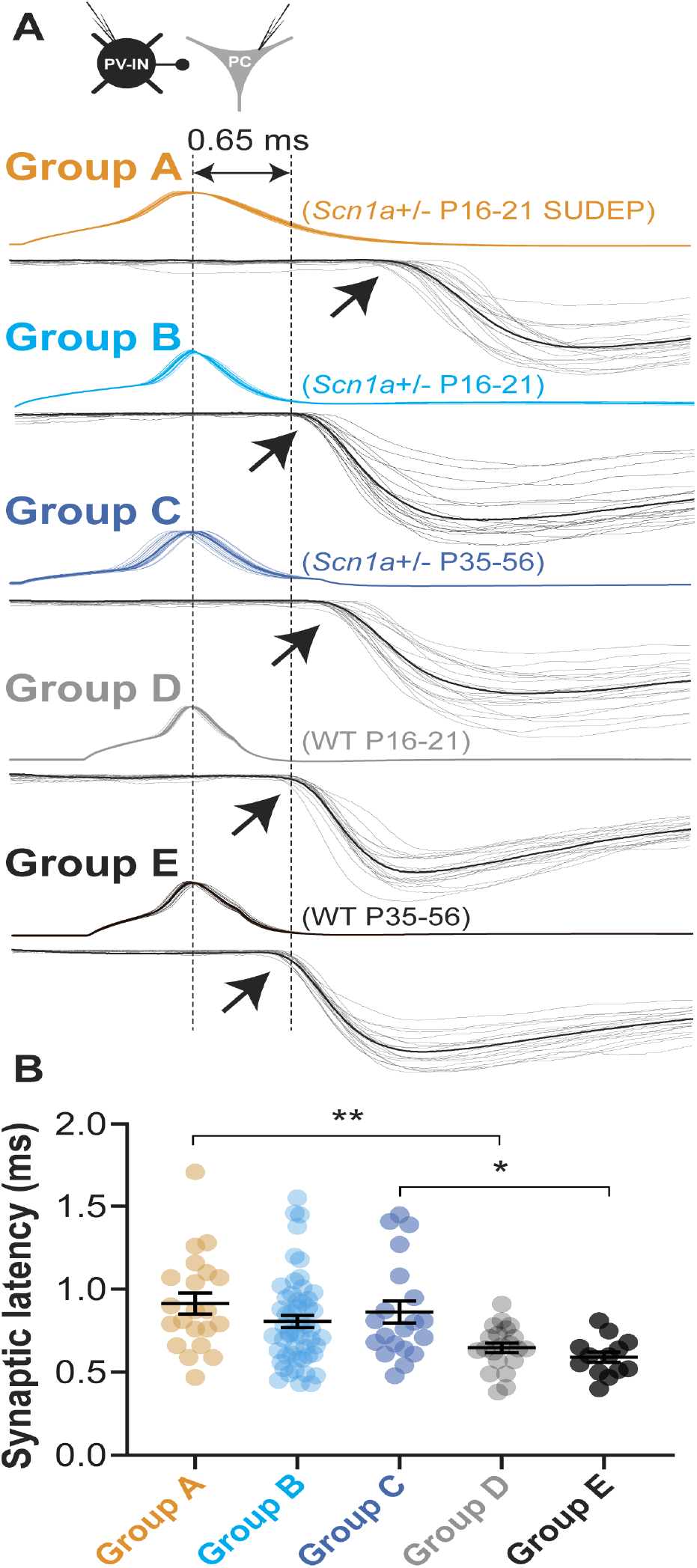
Prolonged synaptic latency at PV interneuron connections. (A) Representative examples from each group illustrating synaptic latency, measured from peak of the presynaptic AP (dotted line at left) to onset (arrows) of the postsynaptic uIPSC (mean, black; individual traces, gray) in Group A-E. The dotted line at right corresponds to a latency of 0.65 ms, added for clarity to approximate the mean value for wild-type (Group D and E). (B) Summary data for synaptic latency in each group. See Table S4.

Synaptic latency is determined by axon length, conduction velocity, and synaptic delay [Sabatini and Regehr, 1999], as well as presynaptic action potential waveform and release probability [Boudkkazi et al., 2007, 2011], and prolonged latency could reflect slowed conduction velocity along PV-IN axons. This interpretation is supported by prior work showing increased synaptic latency at PV-IN:PC synapses in rat dentate gyrus with low concentrations of TTX [Hu and Jonas, 2014]. We and performed multiple whole cell recordings from PV-INs and principle cells in acute brain slices from pre-weanling WT mice (Group D) and found that bath application of low concentrations of TTX (10 nM; estimated to block approximately 50% of Nav1.1-containing voltage-gated Na+ channels; [Smith and Goldin, 1998] had minimal effect on the PV-IN firing pattern (Figure S7A) but led to a two-fold increase in failure rate in a frequency dependent manner similar to what was observed in age-matched pre-weanling P16-21 Scn1a+/− mice (Group A). Failure rate for the fifth pulse in the train was 0.08 ± 0.03 for control and 0.15 ± 0.05 after application of 10 nM TTX (n = 11 connections from 4 mice; p = 0.020 vs. control via two-tailed Student’s t-test); failure rate for the last pulse in a 120 Hz train is 0.12 ± 0.04 in control and 0.23 ± 0.07 after 10 nM TTX (p = 0.047 vs. control). Synaptic latency also increased from 0.49 ± 0.02 to 0.64 ± 0.06 ms after application of TTX (p = 0.016 vs. control). This data supports the finding in Scn1a+/− mice that loss of Nav1.1 in PV-INs may preferentially affect synaptic transmission over spike generation.

### Expression of Nav1.1 at PV Interneuron axons

Direct whole-cell voltage-clamp recordings from PV-IN axons in dentate gyrus indicate that Na+ conductance density increases along the axon [Hu and Jonas, 2014, Hu et al., 2018], suggesting expression of Na+ channels beyond the AIS. Recent work indicates in fact that PV-IN axons in cerebral cortex of rodent and human are extensively myelinated along internodal regions and between axonal branch points [Micheva et al., 2016, Stedehouder et al., 2017, Benamer et al., 2020]. We assessed whether Nav1.1 itself was expressed at PV-IN axons beyond the AIS via immunohistochemistry and confocal microscopy. We confirmed that Nav1.1 is expressed at the AIS of PV-INs (Figure S8A), consistent with previous results [Ogiwara et al., 2007, Li et al., Goff and Goldberg, 2019]. However, we also found Nav1.1 expression along axons, next to branch points (Figure S8B), and at presumptive pre-terminal axons (Figure S8C). We then compared the pattern of Nav1.1 expression to that of pan-Na+ channel labeling (pan-Nav) using automated filtering, segmentation, and extraction from reconstructed Z-stacks (Figure S9A-C) and found that Nav1.1 more prominently labeled branching structures (mean, 3.60 ± 0.24 branches per labeled structure; n = 6,718 puncta from n = 10 Z-stacks from n = 3 mice from young adult wild-type mice) relative to pan-Nav (1.74 ± 0.37 branches per labeled structure; n = 1,409 puncta from n = 7 z-stacks; p < 0.0001 via mixed-effects model) (Figure S9D) (see METHODS). Nav1.1-immunopositive structures were shorter in length (Nav1.1, 33.8 ± 0.92 µm; pan-Nav, 48.5 ± 4.4 µm; p = 0.0009) and were less linear (ratio index for Nav1.1, 2.74 ± 0.03; pan-Nav, 4.98 ± 0.17; p < 0.00001) (Figure S9E-F). We then compared Nav1.1 immunolabeling in young adult WT (Group E) and Scn1a+/− (Group C) mice. We found a marked reduction in the overall normalized intensity of Nav1.1 immunoreactivity in Scn1a+/− relative to wild-type (WT, 96.3 ± 10.6; Scn1a+/−, 72.7 ± 6.0; p = 0.025), with a reduction in the labeling of branching structures (WT, 3.60 ± 0.24 branches/puncta; Scn1a+/−, 2.56 ± 0.10; p = 0.00009) and decreased length of labeled puncta (WT, 33.8 ± 0.92 µm; Scn1a+/−, 29.3 ± 0.3 µm; p < 0.00001) but no change in linearity of puncta (WT, 2.74 ± 0.03; Scn1a+/−, 2.84 ± 0.06; p = 0.013), supporting the possibility that there is a preferential decrease in Nav1.1 at distal axons and branching structures relative to the AIS.

### Alterations in PV Interneuron:Principal Cell Connection Probability

We next assessed for differences in connection rate between PV-INs and PCs in Scn1a+/− vs. wild-type mice, as evidence of decreased connectivity – perhaps due to fewer connections between PV-INs and PCs in Scn1a+/− mice and/or fewer synaptic boutons per synapse at PV-IN:PC connections – could potentially explain the decreased uIPSC amplitude observed in pre-weanling Scn1a+/− mice. The connection rate of neocortical PV-INs to PCs obtained via paired recordings in acute brain slice varies widely, ranging from 25-50% in the published literature [Packer and Yuste, 2011, Espinoza et al., 2018, Vickers et al., 2018], as this depends on slice preparation technique and other considerations (see METHODS). PV-IN:PC synapse formation is also known to change with experience in an activity-dependent manner [Chattopadhyaya et al., 2004], supporting the possibility that this parameter could be altered in Scn1a+/− mice.

We found that PV-INs:PC connection rate was 21.9% (21 connections out of a maximum possible of 96) in pre-weanling Scn1a+/− mice at P16-21 that subsequently were deceased prior to P35 (Group A), yet was 32.6% (105/322) in those that survived to young adulthood (Group B), similar to the connection rate of 35.0% (77/220) observed in age-matched wild-type littermates (Group D; p = 0.023 vs. Group A via chi-square test). Connection rate was 18.8% (46/245) in surviving young adult Scn1a+/− mice at P35-56 (Group C) and 33.8% (52/154) in age-matched wild-type (Group E; p < 0.0011 vs. Group C via chi-square test) (Figure S10B). Hence, we found lower connection probability selectively in pre-weanling Scn1a+/− mice that subsequently were deceased from SUDEP and in surviving young adult Scn1a+/− mice.

Decreased frequency of functional connections between PV-INs and PCs specifically in Groups A and C suggests the possibility that there might be an anatomical difference in synaptic connectivity between groups. To investigate this possibility, we performed immunohistochemical labeling (see METHODS) for synaptotagmin-2 (Syt2), a specific marker of PV-IN synapses (Sommeijer and Levelt, 2012; Bouhours et al., 2017; van Lier et al., 2020), and confirmed that Syt2-positive boutons were perisomatic and PV-positive (Figure S10A). We then quantified the number of PV-IN boutons surrounding the cell bodies of PCs in all experimental groups (Figure S10B). We found a small but statistically significant decrease in the number of Syt2-positive puncta around PCs in Group A relative to Group B and D and between Group C and E (Figure S10C), consistent with a relatively decreased number of synaptic terminals at PV-IN:PC synapses in Groups A and C and consistent with the observed differences in connection probability. The number of perisomatic puncta in layer 4 PCs was 10.4 ± 0.3 in Group A (n = 142 cells from 6 slices from 2 mice), 12.9 ± 0.3 in Group B (n = 147 cells from 8 slices from 6 mice; ANOVA (4, 706) = 1.615; p < 0.0001 vs. Group A with post-hoc correction for multiple comparisons with Holm-Sidak test), 12.7 ± 0.3 in Group C (n = 163 cells from 8 slices from 4 mice; p = 0.65 vs. Group B), 13.3 ± 0.3 in Group D (n = 122 cells from 6 slices from 3 mice; p < 0.0001 vs. Group A; p = 0.65 vs. Group B), and 14.2 ± 0.4 in Group E (n = 134 cells from 7 slices from 3 mice; p = 0.001 vs. Group C; p = 0.12 vs. Group D). Such data indicates a small reduction in perisomatic PV-IN synapses in pre-weanling Scn1a+/− mice at P16-21 deceased prior to P35 (Group A) relative to young Scn1a+/− mice that survive to P35 or later (Group B) and to age-matched WT littermates (Group D), as well as a reduction in perisomatic PV-IN synapses in surviving Scn1a+/− mice at P35-56 (Group C) relative to age-matched young adult WT mice (Group E). These findings may contribute to the observed differences in uIPSC amplitude at PV-IN:PC synapses in Scn1a+/− mice but cannot account for the observed increases in synaptic failure rates.

### A Multicompartment Interneuron Model Supports a Preferential Impact of Reduction in Na+ Conductance on Action Potential Propagation

Because we could not directly confirm that the cause of synaptic transmission failure was due to failure of action potential propagation, we constructed a multi-compartment model of a neocortical PV-IN to assess the differential effects of reduction of Na+ current density at the soma/AIS vs. axon on features of action potential initiation and propagation (see METHODS). Selectively deleting Nav1.1 by 50 % (to model heterozygous loss of Nav1.1 in DS) at specific neuronal compartments (“deletion paradigm”) caused action potential failure (Figure 5). A failure could be that of initiation, or that of propagation (whereby an action potential was initiated at the AIS but not detected at the pre-synaptic terminal) (Figure 5D). When Nav1.1 was partially deleted in the soma, AIS, and Nodes (“—Soma —AIS —Nodes” in Figure 5B-D, modeling Group A), the neuron entered depolarization block at the soma/AIS as seen experimentally in Figure 2. This was also true for the “—Soma —AIS” deletion paradigm. However, when Nav1.1 was impaired specifically at the nodes (“—Nodes” deletion paradigm, modeling Group C), action potentials continued to be generated at the AIS, but were not propagated to the pre-synaptic terminal. The firing rate for both AIS and pre-synaptic terminal recording sites dramatically decreased with further Nav1.1 deletion in both the “—Soma —AIS —Nodes” and “—Soma —AIS” paradigms. Progressive deletion at the nodes only led to a substantial drop-off in firing rate at the presynaptic terminal, with no effect on the firing rate at the AIS.

**Figure 5:**
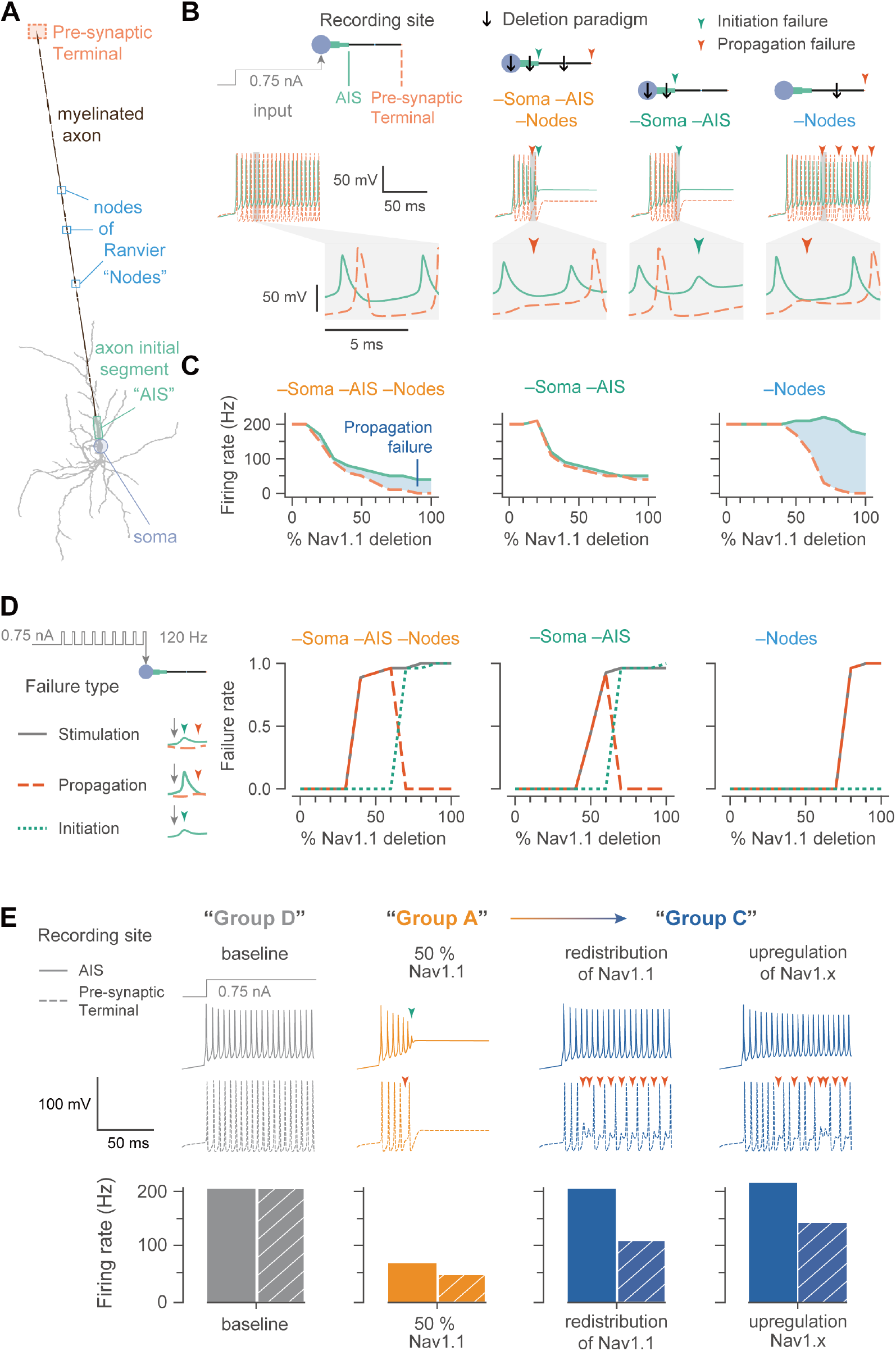
A multi-compartment model supports the selective effect of Na+ reduction at the axon on the fidelity of synaptic transmission at an interneuron synapse. (A) A morphologically realistic neocortical parvalbumin-positive interneuron with a single long myelinated axon interspersed with nodes of Ranvier (“Nodes”), the last of which is the pre-synaptic terminal. (B) Shown are traces of the membrane potential recorded at the axon initial segment (AIS; solid light green line) and presynaptic terminal (dashed light orange line), following injection of 0.75 nA into the soma starting at t = 20 ms. The enlarged trace shows the typical progression of an action potential generated at the AIS and propagated along the axon to the pre-synaptic terminal. The compartment of Nav1.1 deletion is indicated by a black downward arrow, the combination of which is a “deletion paradigm”. Colored downward arrows represent the time of a failure (initiation, dark green; propagation, dark orange), the expected time that an action potential would occur. (C) The average firing rate of a neuron with progressive Nav1.1 deletion. The difference in firing rates between the AIS (solid light green line) and the pre-synaptic terminal (dashed light orange line) represents the propagation failure (shaded blue area), which increases with further deletion. (D) Given current injected at the soma at 120 Hz, the interneuron could exhibit different types of action potential failures similar to ‘B’. First, a failure of stimulation was where a pulse was delivered but an action potential was not detected at the pre-synaptic terminal within 2 ms. Second, a failure of propagation was where an action potential was generated at the AIS but did not reach the pre-synaptic terminal. Finally, a failure of initiation was considered to occur under conditions when a pulse was delivered, but an action potential was not generated at the AIS. Note that, for example, propagation failure in the –Soma –AIS –Nodes deletion paradigm first appears at 30% Nav1.1 deletion, but then returns to zero at 70% Nav1.1 deletion because there is no spike initiation at this level of Nav1.1 deletion. (E) The model was used to mimic and find explainable parameters for the experimental Groups D, A, and C. Traces were from the AIS (solid lines) and pre-synaptic terminal (dashed lines) with failures represented as in ‘B’. Bar plots are the firing rate of the AIS (solid) and pre-synaptic terminal (hatched pattern) of the traces for quantitative comparison. Baseline Nav1.1 in the model represents Group D in the experimental setup (gray) while 50 % Nav1.1 (in the soma, AIS, and nodes) represents experimental Groups A (orange) or B as there was a noticeable reduction in activity. The model can exhibit two separate cases that match activity as seen in Group C (blue), whereby action potential generation was recovered but action potential propagation was still impaired. The first is a redistribution of Nav1.1 whereby the nodes had much less Nav1.1 than the soma and AIS. Here, the Nav1.1 conductance in soma and AIS were 120 % of their baselines (20 % Nav1.1 addition) and nodal Nav1.1 conductance was 40 % of its baseline (60 % Nav1.1 deletion). The second approach to qualitatively mimic Group C activity was to upregulate another transient sodium channel, such as Nav1.3 or Nav1.6. Here, the transient sodium channel was upregulated by 120 % in the soma and 130% in the AIS. See supplementary material for details. See METHODS for details.

We then stimulated the model with brief repetitive depolarizing current injections (Figure 5D). In this model, a failure could be that of stimulation (a pulse was sent but an action potential was not recorded at the pre-synaptic terminal within 5 ms), initiation (a pulse was sent but an action potential was not recorded at the AIS), or propagation (a pulse was sent and an action potential was recorded at the AIS, but not at the pre-synaptic terminal). The progressive loss of Nav1.1 in the “–Soma –AIS –Nodes” deletion paradigm yielded substantial propagation failures that was first apparent at 40% Nav1.1 deletion, and initiation failures at 30% Nav1.1 deletion. Similarly, the “–Soma –AIS” deletion paradigm exhibited propagation failures from 50% Nav1.1 deletion and initiation failures from 70% Nav1.1 deletion. As expected, the “– Nodes” deletion paradigm showed only propagation failures, which were apparent from 80% Nav1.1 deletion.

The model was also used to explore possible mechanisms for the functional recovery seen in Group C (persistent impairment of synaptic transmission with selective recovery of action potential generation) (Figure 5E). One option explored was the possibility that PV-INs might have a fixed pool of Nav1.1 protein that was redistributed from the axon to the AIS, leading to recovery of normal spike generation but worsening of spike propagation. The second option involved upregulation of another transient Na+ conductance, Nav1.X, modeling a condition of compensatory upregulation of, for example, Nav1.3 and/oror Nav1.6 subunits, as has been proposed [Favero et al., 2018] (Figure S11).

The results of the model provide proof of principle that Nav1.1 deletion in specific compartments leads to differential effects on action potential initiation and propagation, and supports the mechanism of transition from Group A to Group C as potentially occurring via redistribution of Nav1.1 or the upregulation of other non Nav1.X Na+ channel subunits.

### A Dual Optogenetic Approach Supports Existence of Frequency-dependent Spike Propagation Failure in Scn1a+/− Mice

Results indicate that PV-IN spike generation is impaired at P16-21 in Scn1a+/− mice but normalizes by P35-56. However, in those mice that survived to P35-56, we found prominent activity-dependent failure of spike propagation, suggesting that, at this time point, the somatically-generated action potential does not reliably propagate to the synapse and trigger neurotransmitter release. To confirm this inference and further compare the fidelity of somatically-generated spikes in wild-type vs. Scn1a+/− mice, we used a dual optogenetic approach in which an excitatory opsin was expressed selectively at the PV-IN soma while a second spectrally distinct opsin was expressed preferentially at synaptic terminals (Figure 6A and Figure S12), facilitating selective generation of somatic vs. axo-synaptic spikes. We used Cre-dependent AAVs to express the excitatory opsin hChR2 (AAV9.Ef1a.DIO.hChR2(E123A).EYFP), which is preferentially targeted to the axon and synapse [Grubb and Burrone, 2010, Mattis et al., 2012, Baker et al., 2016], as well as a soma-targeted red-shifted opsin (AAV9.hSyn.DIO.ChrimsonR.mRuby2.ST; Pégard et al. [2017]), in Scn1a.PV-Cre mice and wild-type PV-Cre littermate controls. We then performed whole-cell voltage-clamp recording of PCs, and recorded IPSCs in response to blue (490 nm; ChR2) and red-light (660 nm; ChrimsonR) photostimulation (optogenetically-evoked IPSCs, or oIPSCs) (Figure 6C-D). We used the mini-slice technique to then compare Scn1a+/− (Groups A-B) and WT (Group D) at P16-21, as well as mutant (Group C) and WT (Group E) at P35-56. Immunohistochemistry and confocal microscopy confirmed that ChrimsonR was selectively expressed at the soma of PV-INs, and hChR2 was expressed preferentially long the axon and at synaptic terminals (Figure 6B).

**Figure 6:**
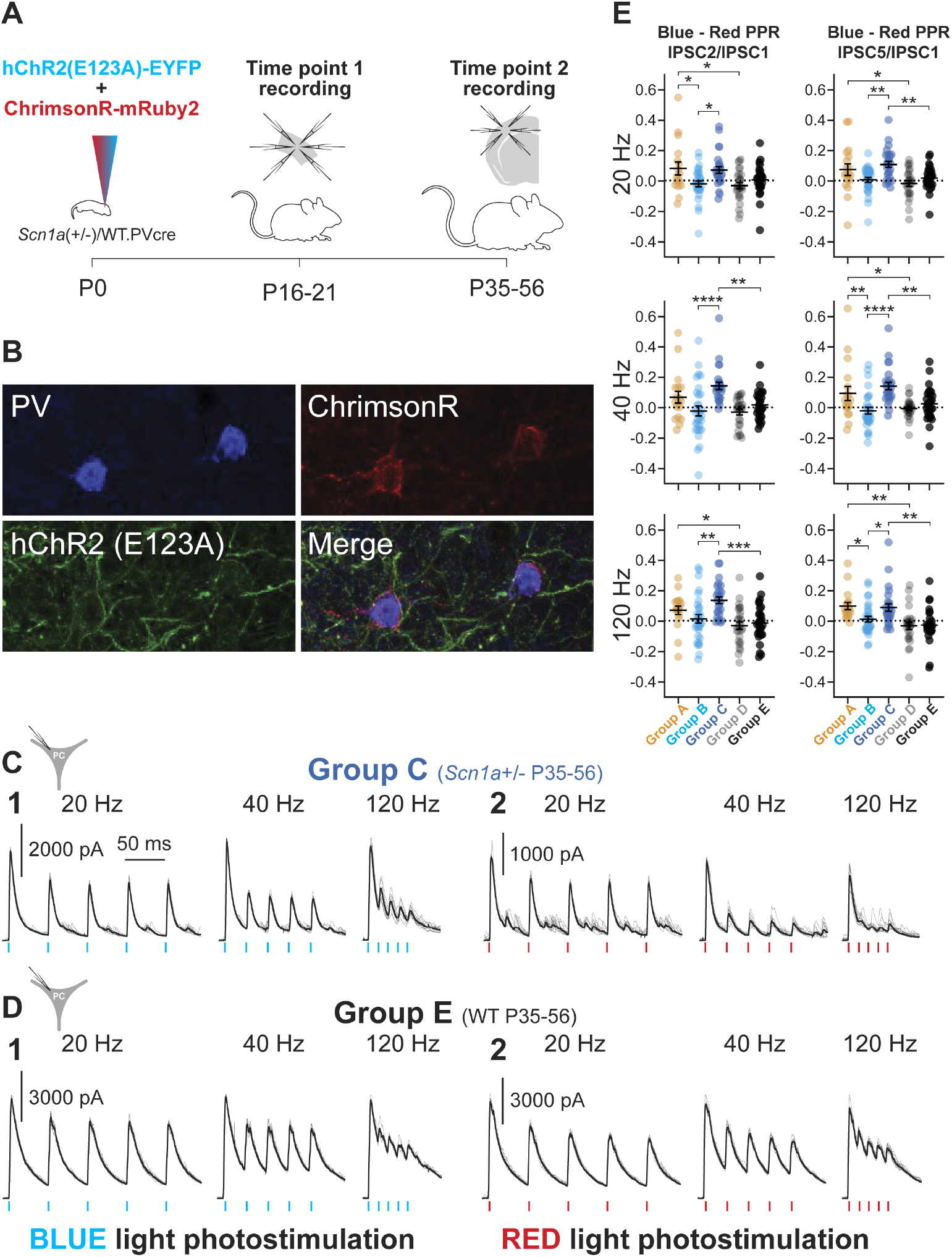
A dual optogenetics strategy demonstrates impaired PV interneuron spike propagation in Scn1a+/− mice. (A) Timeline showing subdural viral injection of both AAV9.DIO.ChrimsonR.mRuby2 and AAV.DIO.hChR2.eYFP into PV-Cre mice at P0 (left) and time points 1 (mini-slice; center) and 2 (acute brain slice; right) recordings. (B) Representative confocal images showing expression of PV (Alexa 405; blue), hChR2.eYFP; green), and ChrimsonR.mRuby2 (red). (C-D) Representative data for evoked oIPSCs recorded in principal cells in response to repetitive photostimulation at 20, 40, and 120 Hz in Group C (C) and E (D). hChR2 was photostimulated with 1 ms pulses at 490 nm (C1 and D1) while ChrimsonR was photoactivated with 1 ms pulses at 660 nm (C2 and D2). (E) Summary data for the index of oIPSCn/oIPSC1 in response to blue light photostimulation minus oIPSCn/oIPSC1 in response to red light photostimulation (Blue – Red index). See Table S5, Table S6, and Table S7.

To isolate oIPSCs, we used the cesium methanesulfonate-based, low chloride-containing internal solution (E_Cl_*−* = -110 mV), with oIPSCs recorded as large outward currents at a holding potential of +10 mV (see METHODS). Electrophysiological recording of PCs combined with photostimulation of surrounding PV-INs revealed a progressive decrease in the amplitude of successive oIPSCs with repeated red light (somatic) simulation that was far greater than that observed in response to blue light (axon/terminal) stimulation (Figure 6C-E). We quantified the ratio of oIPSC2/oIPSC1 and oIPSC5/oIPSC1 at 20, 40 and 120 Hz for blue light (hChR2) and red light (ChrimsonR) stimulation (Figure 6C and Figure 6D, Figure S12, Table S5, and Table S6). To compare responses between groups, we calculated a blue:red index, defined as the arithmetic difference between the blue oIPSC2/oIPSC1 or oIPSC5/oIPSC1 ratio and the red oIPSC2/oIPSC1 or oIPSC5/oIPSC1 ratio. A value of zero reflects no difference in the dynamics of the response to red and blue light and suggests and the somatically-generated response is equally efficacious to the axosynaptic response; this index yields a positive value if there is a greater decrease in oIPSC amplitude with successive stimulation in response to red (somatic) relative to blue light and reflects enhanced efficacy of the axosynaptic response during repetitive stimulation (Figure 6E and Table S7).

There was no significant difference between genotypes or ages in the axon/synaptic terminal evoked oIPSC2/oIPSC1 and oIPSC5/oIPSC1 ratio. However, we found marked differences in soma-evoked oIPSC5/oIPSC1 and oIPSC2/oIPSC1 ratio between groups (Figure S12, Table S5, and Table S6). This yielded a blue:red index value near zero for Groups B, D, and E, indicating no difference in the frequency-dependent dynamics of the oIPSC between somatic and axosynaptic photostimulation. However, we found that Groups A and C exhibited significantly larger values for this blue:red index (Figure 6E and Table**??**), indicating a relative reduction in the somatically-evoked oIPSC response. For example, the oIPSC5/oIPSC1 index at 40 Hz was 0.10 ± 0.05 for Group A (n = 18 PCs from 6 mice), - 0.02 ± 0.02 for Group B (n = 30 PCs from 11 mice; ANOVA F(4, 123) = 2.137; p = 0.0098 vs. Group A with post-hoc correction for multiple comparisons with Holm-Sidak test), 0.15 ± 0.03 for Group C (n = 26 PCs from 7 mice; p < 0.0001 vs. Group B), 0.00 ± 0.00 for Group D (n = 26 PCs from 8 mice; p = 0.0289 vs. Group A; p = 0.6654 vs. Group B), and 0.0300 ± 0.02 for Group E (n = 28 PCs from 8 mice; p = 0.0031 vs. Group C and 0.6034 vs. Group D). Using targeted current clamp recordings, we confirmed that PV-INs at all age groups could reliably follow brief (1 ms) pulses of red light photostimulation at 120 Hz (Figure S12).

This result is consistent with a frequency- and activity-dependent failure of synaptic transmission with somatic stimulation that can be “bypassed” via direct axo-synaptic stimulation. Such findings support the results of the multiple whole-cell electrophysiology experiments indicating that failures of synaptic transmission reflect frequency-dependent failure of the somatic action potential to drive neurotransmitter release.

## DISCUSSION

Major advances in genetic diagnostic testing over the last twenty years has led to the identification of hundreds of genes associated with neurodevelopmental disorders [Parenti et al., 2020], including intellectual disability, ASD, and epileptic encephalopathy. Understanding how such genetic variation leads to disease has proven challenging [Sahin and Sur, 2015, del Pino et al., 2018]. This remains the case for Dravet syndrome, despite being the most common of the epileptic encephalopathies, affecting approximately 1 in 15,000 individuals in the United States [Wu et al., 2015]; SCN1A is in turn the most frequently identified epilepsy gene, with pathogenic variants in SCN1A associated with an estimated 0.2% of all epilepsies [Meng et al., 2015, Symonds et al., 2019].

Our prior work demonstrated that abnormalities in PV-IN spike generation and repetitive action potential discharge identified in Scn1a+/− mice was in fact transient and restricted to early (pre-weanling) developmental time points (second and early third postnatal weeks) [Yu et al., 2006, Ogiwara et al., 2007, Tai et al., 2014, Tsai et al., 2015, De Stasi et al., 2016]; PV-INs recorded in young adult mice (P35-56) had seemingly recovered the ability to discharge sustained trains of action potentials at high frequency [Favero et al., 2018]. We showed that the axon initial segment – defined by ankyrin-G immunoreactivity – was longer in PV-INs from Scn1a+/− mice as compared to age-matched WT controls (perhaps suggesting a prior history of decreased excitability; [Grubb et al., 2011, Bender and Trussell, 2012, Wefelmeyer et al., 2016, Yamada and Kuba, 2016]). In addition, we had found a clear “hump” in the dV/dt plot in mature (P35+) PV-INs in Scn1a+/− mice that was not seen in WT PV-INs – a finding confirmed in the present study (Figure S4A) – suggesting spatiotemporal separation between the AIS spike and somatic spike, which we hypothesized to be due to an extension of or distal shift in the location and/or extent of the AIS perhaps secondary to upregulation of non-Nav1.1 Nav1.X Na+ channel subunits, although this could also be due at least in part to slowing of spike propagation.

However, Scn1a+/− mice in the chronic phase continue to exhibit seizures (albeit at a decreased frequency), ASD-related endophenotypes such as impaired social interaction and novel object recognition, and SUDEP-like phenomena – as is the case with human patients – and which cannot be explained by impaired action potential generation by neocortical PV-INs. Hence, the primary goal of the present study was to investigate PV-IN synaptic transmission in Scn1a+/− mice as an indirect output measure of spike propagation. We hypothesized that PV-INs are able to reconstitute high-frequency firing at the soma via unidentified compensatory plasticity mechanisms, yet cannot recover axonal excitability resulting from heterozygous loss of Nav1.1. We confirmed that PV-INs in young adult Scn1a+/− mice (Group C) indeed recover fast spiking, yet exhibit impaired synaptic transmission as evidenced by markedly increased synaptic failure rates (Figure 3) and prolonged synaptic delay (Figure 4) at PV-IN:PC synapses. This suggests that PV-IN axonal dysfunction is an important locus of pathology in the chronic phase of Dravet syndrome. Such a conclusion makes logical sense as prior work has shown that low concentrations of TTX estimated to block 50% of voltage-gated Na+ channels produce only mild effects on action potential generation in PV-INs, consistent with the presence of a “supercritical” density of Na+ channels at the PV-IN AIS (Hu and Jonas, 2014) and a relatively higher safety factor for spike generation at the AIS. However, such a high density of Na+ channels is required for spike reliability and propagation speed. While PV-INs in pre-weanling Scn1a+/− mice do exhibit impairments in action potential generation, these cells are still able to discharge sustained trains of action potentials at > 200 Hz, just as PV-INs qualitatively retain fast-spiking capabilities in the presence of low concentrations of TTX. Our computational model supports this conclusion, demonstrating selective effects of reduced Nav1.1 conductance on the fidelity of spike propagation relative to spike initiation in a detailed multicompartment model of a neocortical PV-IN informed by detailed biological data on somatic and axonal conductances and axonal anatomy.

It should be noted that there is unlikely to be a singular cellular/circuit mechanism of DS, or of any complex neurodevelopmental disorder. It is known that excitability of SST and VIP-INs are also impaired in DS (Tai et al., 2014; Rubinstein et al., 2015; Almog et al., 2021), and that VIP-IN dysfunction persists in young adult Scn1a+/− mice (Goff and Goldberg, 2019). This work focused on neocortical PV-INs, yet it is clear that the hippocampus and, in particular, the dentate gyrus, is a prominent locus of pathology in DS models [Liautard et al., 2013, Dutton et al., 2017, Stein et al., 2019, Mattis et al., 2021]and in human patients [Van Poppel et al., 2012, Gaily et al., 2013].

### The Mini-Slice Technique to Evaluate Cell-Type Specific Electrophysiological Properties Across Development

Our prior work [Favero et al., 2018, Tran et al., 2020, Mattis et al., 2021] and much of the work in the field was limited by the obvious fact that we could only record from young adult Scn1a+/− mice from those mice that had actually survived. Hence, it could be the case that PV-INs found to exhibit normal high-frequency firing at P35+ had always been normal, with PV-IN dysfunction being present only in those mice decreased prior to P35. Hence, a secondary aim of the study was to resolve this issue, which required development of a method to perform targeted recordings from PV-INs from the same animal at two different (early, P16-21; late, P35-56) time points, so as to allow for within-animal comparison. This novel approach confirmed that PV-IN dysfunction is preset at early time points even in those animals that survive to young adulthood, and allowed us to show that recovery of PV-IN action potential generation occurs in the same animals in which PV-IN excitability was directly shown to be dysfunctional at earlier time points. Furthermore, the identification of combined abnormalities in spike generation and synaptic transmission in Scn1a+/− mice that exhibit early mortality may represent a marker predictive of SUDEP that could be targeted for modification. This mini-slice technique could prove widely applicable to the study of risk factors for mortality in preclinical models of severe neurological disease.

### Combined Early Impairments in Spike Generation and Spike Propagation may be a Risk Factor for Early Mortality

PV-INs from all Scn1a+/− mice showed impairments in spike generation at early time points regardless of survival to or death prior to P35, while PV-INs had recovered high-frequency firing in Scn1a+/− mice surviving to adulthood. However, the fidelity of inhibitory synaptic transmission at PV-IN:PC synapses was decreased selectively at P16-21 in those Scn1a+/− mice that subsequently died of SUDEP (Group A) relative to mice that survived (Group B) (Figure 3C and Figure 3D). This was confirmed by a dual optogenetics approach that allowed us to separate somatic and axo-synaptic stimulation of PV-INs (Figure S11), and revealed a relative impairment of somatically-generated but not axo-synaptic oIPSCs (Figure 5E), suggesting that direct stimulation of PV-IN terminals can bypass PV-IN axonal dysfunction. And while PV-INs from young adult Scn1a+/− mice (Group C) did not show impairments in spike generation (Figure 2, Figure S1C, Figure S2, and Figure S4; consistent with the results of our previous study; Favero et al. [2018]), we found frequency-dependent impairment in synaptic transmission at PV-IN:PC synapses in Group C, similar to that observed in Group A (Figure 3, Figure 4, Figure 5). These results suggest that: (1) combined early impairments in spike generation and synaptic transmission may be a risk factor for early mortality in Scn1a+/− mice; (2) severe impairments in synaptic transmission are involved in the pathophysiology of continued symptoms of Dravet syndrome in the chronic phase; and (3) targeted modulation of axonal or presynaptic function may represent a therapeutic approach to this disorder.

### Implications for Development of Novel Therapies for Dravet Syndrome

Here we provide the first evidence that PV-INs in Scn1a+/− mice exhibit impaired synaptic transmission, that early impairments in PV-IN synaptic transmission may predict SUDEP risk, and that such deficits persist across developmental time and may contribute to the pathology characteristic of Dravet in the chronic phase of the disorder. This work has important implications for emerging efforts to increase the expression of SCN1A as a potential therapeutic strategy [Colasante et al., 2019, Han et al., 2020, Yamagata et al., 2020]. Upregulation of SCN1A may be ineffective or even deleterious if administered after disease onset if chronic dysfunction is driven by a mechanism other than haploinsufficiency of Nav1.1. Of note, AAV-mediated upregulation of the voltage gated Na+ channel *β*1 accessory subunit – which regulates *α* subunit expression, plasma membrane localization, and subcellular targeting, among other functions [O’Malley and Isom, 2015] – led to decreased seizure frequency, improved performance on cognitive tests, and reduced mortality in Scn1a+/− mice [Niibori et al., 2020]. Our findings suggest that increasing the fidelity of PV-IN synaptic transmission, perhaps via upregulation of Nav1.1 specifically in the axosynaptic compartment such as at nodes of Ranvier and/or axonal branch points of PV-INs [Micheva et al., 2016, Stedehouder et al., 2017, Benamer et al., 2020], or by otherwise bypassing axonal dysfunction, may be important for the treatment of Dravet syndrome-related pathology.

## METHODS

### Key Resource Table

#### Animals

All procedures and experiments were approved by the Institutional Animal Care and Use Committee at the Children’s Hospital of Philadelphia and were conducted in accordance with the ethical guidelines of the National Institutes of Health. Both male and female mice were used in equal proportions. After weaning at P21, mice were group-housed with up to five mice per cage and maintained on a 12-hour light/dark cycle with ad libitum access to food and water.

Mouse strains used in this study included: Scn1a+/− mice on a 129S6.SvEvTac background (RRID: MMRRC_037107-JAX) generated by a targeted deletion of exon 1 of the Scn1a gene, PV-Cre mice (B6;129P2-Pvalbtm1(cre)Arbr/J; RRID:IMSR_JAX:017320); tdTomato reporter/Ai14 mice (Rosa-CAG-LSL-tdTomato; RRID: IMSR_JAX:007914; on a C57BL/6J background); wild-type 129S6.SvEvTac (Taconic Biosciences model #129SVE; RRID:IMSR_TAC: 129sve); and wild-type C57BL/6J (RRID:IMSR_JAX:000664).

Homozygous PV-Cre mice were crossed to homozygous Ai14 mice to generate PV-Cre.tdT double heterozygotes on a C57BL/6J background. Female PV-Cre.tdT double heterozygotes were then crossed to male 129S6.Scn1a+/− mice to generate Scn1a.PV-Cre.tdT mice and WT. PV-Cre.tdT littermate controls. The genotype of all mice was determined via PCR of tail snips obtained at P7 and was re-confirmed for each mouse after they were sacrificed for slice preparation. All mice used for experiments were on a near 50:50 129S6:B6J background, and Scn1a+/− mice on this background have been shown to replicate the core phenotype of Dravet Syndrome [Miller et al., 2014, Mistry et al., 2014]. We observed similar rates of spontaneous death (22/84, or 26.2% by P56, for Scn1a.PV-Cre.tdT mice; [Favero et al., 2018]).

## METHOD DETAILS

### Viral injections

Adeno-associated viruses (AAV) encoding the red-shifted channelrhodopsin variant ChrimsonR (Klapoetke et al., 2014) tagged with mRuby2 (AAV9.hSyn.DIO.ChrimsonR.mRuby2.ST; Addgene Catalog No. 105448; titer, 2.13 X 1013 GC/mL) and eYFP-tagged hChR2 tagged (AAV9.EF1a.DIO.hChR2(E123A).EYFP; Addgene Catalog No. 35507; titer, 0.73 X 1013 GC/mL) were injected into S1 through the skull of mice age P0 using a NanoFil syringe (10 µl; WPI) under ice anesthesia. Mice recovered on a heating pad within 10 min and were placed in their home cage.

### Acute slice preparation

Mice were deeply anesthetized with 5% isoflurane and transcardially perfused with ice cold aCSF containing (in mM): KCl, 2.5; NaH2PO4, 1.25; HEPES, 20; N-Methyl-D-glucamine (NMDG), 93; L-ascorbic acid (sodium salt), 5; thiourea, 2; sodium pyruvate, 3; CaCl2, 0.5; MgSO4, 10; D-glucose, 25; N-acetyl-L-cysteine, 12; NaHCO3, 30; with pH adjusted to 7.30 with HCl and osmolarity adjusted to 310 mOsm, and equilibrated with 95% O2 and 5% CO2. Slices recovered in modified aCSF for 30 min at 32°C. Slices were transferred to standard (recording) aCSF containing (in mM): NaCl, 125; KCl, 2.5; MgSO4, 2; NaH2PO4, 1.25; CaCl2, 2; D-glucose, 20; NaHCO3, 26. Slices were maintained at room temperature for up to 3 hr before recording. Slices were then transferred to a recording chamber on the stage of an upright microscope (Olympus) and continuously perfused using a diaphragm pump (Smoothflow Pump Q Series, Model #Q-100-TT-ULP-ES, TACMINA CORPORATION, Japan) and suction (PC-21, NARISHIGE, Japan) with recording aCSF at a rate of 3 mL/min at 30 ± 1°C.

### Mini-slice preparation

Mice were placed in a stereotaxic apparatus (Kopf Model #963) and deeply anesthetized via inhalation of 3-5% isoflurane in oxygen for induction and maintained at 1.5-2% during the surgery. Under sterile conditions, a small 2.5 X 2.5 mm square craniotomy was made overlying primary somatosensory cortex using a hand-held drill (Osada EXL-M40) equipped a 1.0 mm diameter drill bit (NeoBurr #EF4), and an approximately 2 X 2 X 0.8 mm depth rectangular biopsy specimen was removed with the tip of a #10 scalpel blade (Figure 1Ba). The collected tissue was immediately placed into ice-cold, oxygenated, modified aCSF (see Acute slice preparation) containing 3% low-melting point agarose (Lonza; Catalog Number 50101). The tissue block was allowed to harden in a plastic mold on ice and then mounted immediately on the specimen holder of a Leica VT-1200S vibratome using cyanoacrylate glue and sliced at 300 µm. Hemostasis was achieved with Gelfoam® sterile sponges (Pfizer) soaked in aCSF. The craniotomy bone flap was replaced and the scalp was sutured.

### Recordings of electrophysiological membrane/firing properties and unitary IPSCs

PV-INs were identified by tdTomato expression visualized with epifluorescence. Spiny stellate cells (PCs; principal cells) were identified by morphology under infrared differential interference contrast (IR-DIC) and the presence of a regular-spiking firing pattern. Whole-cell recordings were obtained from layer 4 primary somatosensory cortex (S1; “barrel”). Patch pipettes were pulled from borosilicate glass using a P-97 puller (Sutter Instruments) and filled with intracellular solution containing (in mM): K-gluconate, 65; KCl, 65; MgCl2, 2; HEPES, 10; EGTA, 0.5; Phosphocreatine-Tris2, 10; ATP (magnesium salt), 4; GTP (sodium salt), 0.3; pH was adjusted to 7.30 with KOH, and osmolarity adjusted to 290 mOsm. Pipettes had a resistance of 3-4 MΩ when filled and placed in recording solution. Unitary IPSCs were obtained via 2-6 simultaneous patch clamp recordings from neurons located within 100 µm of one another in a mini-slice in pre-weanling mice (Groups A-C; Figure 1B) or a thalamocortical slice (Agmon and Connors, 1991) in youngadult mice (Groups C and E; Figure 3A1 and Figure 3A2).

Voltage was sampled at 100 kHz with a MultiClamp 700B amplifier (Molecular Devices), filtered at 2 kHz by Lowpass and Bessel filters, digitized using a DigiData 1550B, and acquired using pClamp11 software. Recordings were discarded if the cell had an unstable resting membrane potential and/or a membrane potential of PV-INs greater (less negative) than -60 mV, or if access resistance increased by > 20% during the recording. We did not correct for liquid junction potential.

### Recordings of optogenetic evoked IPSCs

PV-INs were identified by mRuby2 expression visualized with epifluorescence and PCs were identified as above. To record optogenetically-evoked spikes in PV-INs (Figure S10), we used a regular intracellular solution containing (in mM): K-gluconate, 130; KCl, 6.3; MgCl2, 1; HEPES, 10; EGTA, 0.5; Phosphocreatine-Tris2, 10; ATP (magnesium salt), 4; GTP (sodium salt), 0.3; pH was adjusted to 7.30 with KOH, and osmolarity adjusted to 290 mOsm. To record optogenetic evoked IPSCs in PCs (Figure 5), we used a cesium-based intracellular solution containing, in mM: cesium methanesulfonate, 125; HEPES, 15; EGTA, 0.5; QX-314, 2; Phosphocreatine-Tris2, 10; tetraethylammonium chloride, 2; ATP (magnesium salt), 4; GTP (sodium salt), 0.3; pH was adjusted to 7.30 with KOH, and osmolarity adjusted to 290 mOsm.

### Immunohistochemistry

For visualization of PV-IN axonal boutons, microbiopsies collected from young Scn1a+/− mice (Groups A and B) were drop-fixed in 4% PFA. Mice from Groups C, D, and E were deeply anesthetized with isoflurane, then transcardially perfused with 10-15 mL of 4% PFA in PBS and postfixed for 24 h. Brain tissue was then transferred to 30% sucrose solution for 24-48 h and then 40 µm coronal sections through S1 were made using a sliding microtome (American Optical). After washing in PBS, the slices were blocked (3% normal goat serum, 2% bovine serum albumin, 0.3% Triton X-100 in PBS) for 1 h at room temperature to decrease nonspecific staining, and then washed in PBS. The sections were then transferred into solution containing mouse-*α*-synaptotagmin-2 primary antibody (1:5000; ZFIN ZDB-ATB-081002-25) with 1% normal goat serum, 0.2% bovine serum albumin, and 0.3% Triton X-100 in PBS at 4°C overnight in the dark. After washing in PBS, the incubated sections were transferred into a solution containing the secondary antibody (Alexa Fluor 488-conjugated goat anti-mouse IgG2a; 1:500, Molecular Probes A21131) with 2% bovine serum albumin and 0.3% Triton X-100 in PBS at 4°C overnight in the dark. Sections were then mounted on glass slides and imaged using a confocal microscope (Leica SP8) equipped with a 40X objective (Leica HC PL APO 40X NA 1.30 OIL CS2). Bouton density was counted surrounding putative principle cell somata in a randomly selected 100 X 100 µm area in layer 4 primary somatosensory neocortex. For validation of optogenetic experiments, Scn1a.PV-Cre.tdT and WT.PV-Cre.tdT mice injected in the subdural space with AAV9.hSyn.DIO.ChrimsonR.mRuby2.ST and AAV9.EF1a.DIO.hChR2(E123A).EYFP at P0 and processed for immunohistochemistry at P35-56 as described above. Tissue sections were transferred to a solution containing rabbit anti-PV antibody (1:1000; Swant PV27), 1% normal goat serum, 0.2% bovine serum albumin, and 0.3% Triton X-100 in PBS, and incubated at 4°C overnight in the dark. After washing in PBS, the incubated sections were transferred into solution containing secondary antibody (Alexa Fluor 405; 1:500, Thermo Fisher A31556) with 2% bovine serum albumin and 0.3% Triton X-100 in PBS at 4°C overnight in the dark. Sections were mounted on glass slides and confocal microscopy was performed to visualize cells immunopositive for PV and co-expressing hChR2-eYFP and ChrimsonR-mRuby2.

Na+ channel immunohistochemistry was performed on free-floating brain tissue sections following light fixation and permeabilization as described previously [Goff and Goldberg, 2019]. Briefly, isoflurane-anesthetized mice were transcardially perfused with 1% paraformaldehyde and 0.5% MeOH in PBS; then, the brain were removed and post-fixed in perfusate at room temperature for 1 hour. 50 µm thick sections were cut on a Leica VT-1200S vibratome, and then blocked and permeabilized with 0.2% Triton X-100 (Sigma) and 10% normal goat serum in PBS for one hour at room temperature. Slices were stained overnight at 4° C with a primary antibody directed against Nav1.1 (NeuroMab K74/71) or Pan-Nav (NeuroMab N419/40) in PBS with 3% bovine serum albumin (BSA, Sigma) and 0.2% Triton X-100. The following day, the slices were washed with PBS and stained with a secondary antibody, Alexa Fluor 488-conjugated goat anti-mouse IgG1 or IgG2 respectively (Molecular Probes), in PBS, with 3% BSA and 0.2% Triton X-100. Slices were mounted, cover-slipped, and sealed before imaging on a Leica TCS SP8 confocal microscope. All imagining was done at 80X using identical laser power, exposure, and averaging parameters. Several Z-stacks of Layer IV barrel cortex and spanning the thickness of each section were taken using 1 µm steps for each sample. The dimensions of each Z-stack were 150 µm X 150 µm X 20-40 µm thickness.

For automated analysis of confocal images, Z-stacks were first imported into ImageJ, and the green channel (Nav1.1 or Pan-Nav) was exported as a .tif image sequence excluding any sections near the surface of the slice which tended to show brighter non-specific labelling. These sequences were imported into Matlab, where each image was segmented in the following sequence: each plane in the sequence was binarized using Otsu’s method to determine a global threshold; then, the activecontour function was used to refine the generated mask for 100 iterations using the edge detection method; and finally, the image mask was refined using imopen and imfill. The utilized functions are all available through the Matlab Image Processing and computer vision toolbox, and accessed through the image segmenter app. A fraction of the segmented images were checked against a blind scorer and were found to be in good agreement with well-defined Nav1.1 and Pan-Nav puncta. Puncta with a total length smaller than 2 µm or with an area greater than 160 µm2 were filtered out of analysis, as these were not usually reported as staining by a blind scorer but rather seemed to be non-specific labelling of large structures like blood vessels/nuclei and visual noise. The individual images were then reconstructed into a 3D volume, and segmentation analysis was done in 3D using the regionprops function. From this, we extracted the major and minor axis length, number of branch points, and a 2D flattened representation of each puncta for calculation of linearity as the ratio of the major to the minor axis. Mean intensity values for each puncta were normalized by subtracting the median background value of each slice of the Z-stack. “Branches” were defined by the built-in Matlab function as the number of branch points for an individual puncta, with a “branch point” corresponding to a point on the extracted 3D volume where a pixel splits into two or more structures. Linearity was defined as the unit-less ratio of major to minor axis.

### Electrophysiology data analysis

All analysis was performed manually in Clampfit (pCLAMP) or using routines written in Matlab. Resting membrane potential (Vm) was calculated using the average value of a 1 s sweep with no direct current injection. Input resistance (Rm) was calculated using the response to hyperpolarizing current injections near rest using Rm = V/I. AP threshold was calculated as the value at which the derivative of the voltage (dV/dt) first reached 10 mV/ms. Spike height refers to the absolute maximum voltage value of an individual AP, while spike amplitude was calculated as the difference between spike height and AP threshold for a given AP. AP rise time is the time from AP threshold to the peak of the AP. AP half-width (AP 1/2-width) is defined as the width of the AP (in ms) at half-maximal amplitude. AP afterhyperpolarization (AHP) amplitude is calculated as the depth of the afterhyperpolarization (in mV) relative to AP threshold. Unless indicated, all quantification of single spike properties was done using the first AP elicited at rheobase. Rheobase was determined as the minimum current injection that elicited APs using a 600 ms sweep at 25 pA intervals. Maximal instantaneous firing was calculated using the smallest interspike interval (ISI) elicited at near-maximal current injection. Maximal steady-state firing was defined as the maximal mean firing frequency during the last 300 ms of a suprathreshold 600 ms current injection, with a minimum requirement for a spike being an amplitude of 40 mV with a clear AP threshold of dV/dt > 10 mV/ms and height overshooting at least 0 mV. All I-f plots were created using the steady-state firing calculated for each current step, counting failures as 0 for subsequent current steps. The amplitude of the unitary IPSC was calculated as the difference between baseline and peak from the average of 10-20 consecutive sweeps obtained with a 15 second inter-sweep interval to facilitate recovery from short-term synaptic depression. The paired pulse ratio (PPR) was calculated from this data as the ratio of the second (IPSC2) to the first IPSC. At 40 and 120 Hz, the amplitude of the subsequent IPSCs in the train was calculated as the difference between the peak amplitude and the extrapolation of the single exponential fit to the decay of the preceding IPSC. uIPSC latency was measured from the peak of the presynaptic AP to the onset of the IPSC. uIPSC failure was defined as the absence of a transient current greater than 5 pA occurring within 5 ms after the presynaptic AP.

For the contingency analysis, all unitary connections exhibiting IPSC1 failure were pooled from Groups A, B, and C. We defined Type 1 IPSC2 as the IPSC2 following success of IPSC1, and Type 2 IPSC2 as IPSC2 following a failure of IPSC1.

### NEURON model

A detailed interneuron model was derived from that specified in [Berecki et al., 2019]. Data on neuronal morphology and ion channel biophysical properties were from the Blue Brain Project neocortical microcircuit portal (https://portal.bluebrain.epfl.ch/resources/models; [Ramaswamy et al., 2015]), and formulated using formulations Hodgkin-Huxley kinetics. Previously, values were fitted using BluePyOpt to fit the electrophysical characteristics of continuous non-accommodating fast-spiking PV-INs. Here, the model was adapted to explore the effects of altered Nav1.1 conductances in the axon initial segment (AIS) and the nodes of Ranvier (“Nodes”) by spacing a 1 *µ* m node every 33 *µ*m [Stedehouder et al., 2017]. In accordance with existing data, the Nodes contained Nav1.1 and Kv3 channels with soma-to-axon and Na+-to-K+ conductance ratios in-line with [Hu et al., 2014]. Simulations were performed using NEURON and Python at 34°C. Current was injected at the soma and action potentials recorded at 26.5 *µ* m (the length of the AIS) and 1056.5 *µ* m (the pre-synaptic terminal and last node). The current was either continuous (i.e., a rectangular pulse delivered in current-clamp mode) or pulsed at specific frequencies. An action potential was measured if the membrane potential crossed -20 mV. Nav1.1 was selectively deleted at specific neuronal compartments, proportional to the baseline conductance of each compartment.

As in Figure 6, failure rate was the number of failures divided by total stimuli. A failure could be that of stimulation (a pulse was sent but an action potential was not recorded, for any reason, at the pre-synaptic terminal within 5 ms), initiation (a pulse was sent but an action potential was not recorded at the AIS), or propagation (an action potential was recorded at the AIS but not at the pre-synaptic terminal).

To assess potentials mechanisms that could exhibit functional recovery as seen in Group B (action potential generation recovery but still exhibiting impaired propagation), Nav1.1 conductances were either redistributed from the nodes to the AIS, or another transient sodium conductance, imitating a possible sources Nav1.3 or Nav1.6 channels, was upregulated. The necessary amount redistribution and upregulation for the model was determined quantitatively (Figure S9).

### Quantification and statistical analysis

Unless indicated, the six key comparisons were performed using one-way ANOVA with post-hoc Holm-Sidak Test to assess for differences between genotype or age. Statistical significance was defined as p < 0.05 after post-hoc correction with p values reported exactly. All average values are mean ± SEM. Numbers for mice, cells, and synapses, are reported in the Supplemental Tables.

Linear mixed modeling was performed on a subset of data to account for the fact that more than one cell or synaptic connection was recorded from some mice. The model was fitted to the data with mouse as a random factor.

## AUTHOR CONTRIBUTIONS

Conceptualization and Methodology: K.K., C.C., T.P.V., and E.M.G.; Formal Analysis: K.K., C.C., K.M.G., A.S., T.P.V., and E.M.G.; Investigation: K.K. C.C., and K.M.G.; Writing – Original Draft: K.K. and E.M.G; Writing – Review and Editing: K.K., C.C., K.M.G., A.S., T.P.V., and E.M.G.; Funding Acquisition: E.M.G.

## ACKNOWLEDGEMENTS

We would like to thank Bernardo Rudy as well members of the Goldberg Lab for comments on a previous version of the manuscript; Xiaohong Zhang for expert technical support and mouse colony maintenance; Jennifer Kearney for the gift of Scn1a+/− mice; and Edward Boyden for the distribution of ChrimsonR and Hillel Adesnik for the distribution of ChrimsonR.mRuby2.ST. This work was supported by the National Institutes of Neurological Disorders and Stroke of the National Institutes of Health under F31NS111803 (to K.M.G.), K08NS097633 and R01NS110869 (to E.M.G), the Dravet Syndrome Foundation (to A.S.), an ERC Consolidator Grant (SYNAPSEEK) (to T.P.V.), and the NOMIS Foundation through the NOMIS Fellowships at IST Austria program (to C.C.).

## SUPPLEMENTAL INFORMATION

Supplemental Information appears below.

---

**Figure S1:**
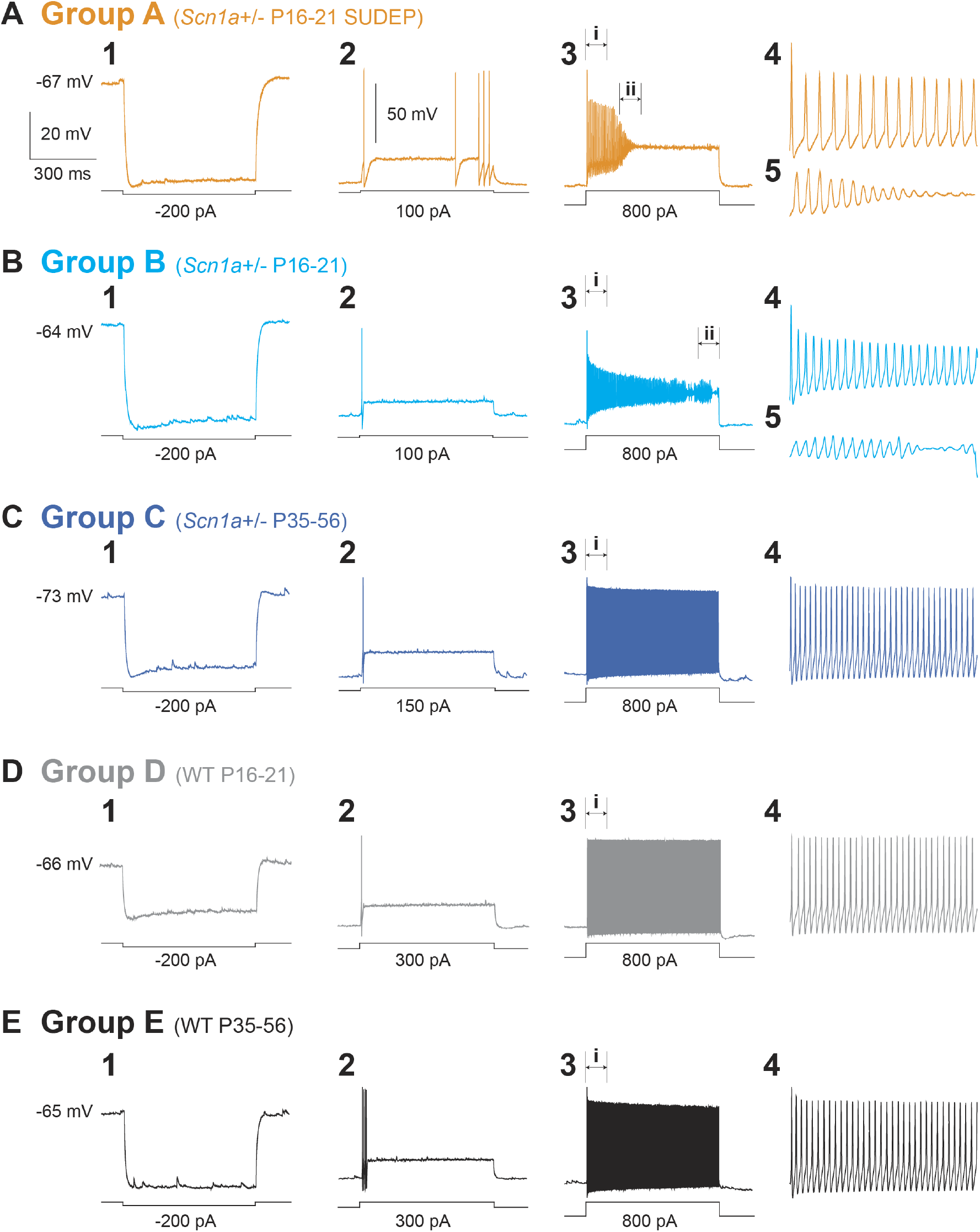
Excitability of PV interneruons in Scn1a+/− vs. wild-type mice. Representative traces of PV-INs from Group A (A), B (B), C (C), D (D), and E (E). Shown is the response to a -200 pA hyperpolarizing current injection (A1-E1), a near-rheobase current injection (A2-E2), and an 800 pA depolarizing current injection (A3-E3), with an expanded view at onset of firing (A4-E4) and at spike failure (A5 and B5).

**Figure S2:**
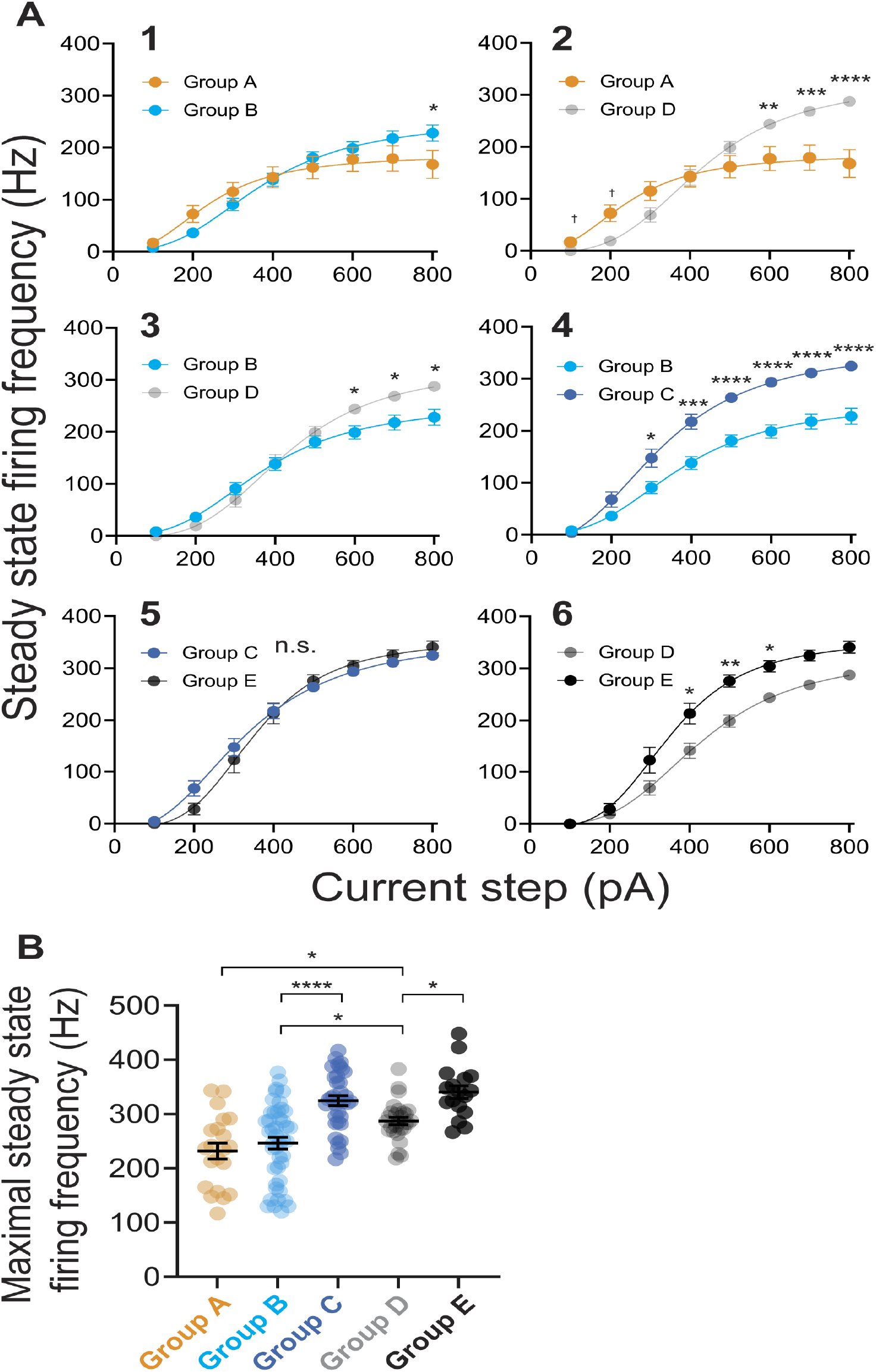
Decreased maximal steady-state firing frequency in PV interneurons from Scn1a+/− mice at early developmental time points. (A) I–f curves for the six key inter-Group Comparisons showing the relationship between current injection and maximal steady state firing frequency, from Figure 2B. Group A vs. Group B (1), Group A vs. Group D (2), Group B vs. Group D (3), Group B vs. Group C (4), Group C vs. Group E (5), and Group D vs. Group E (6). See also Table S2. (B) Summary data for maximal steady state firing frequency. See also Table S2.

**Figure S3:**
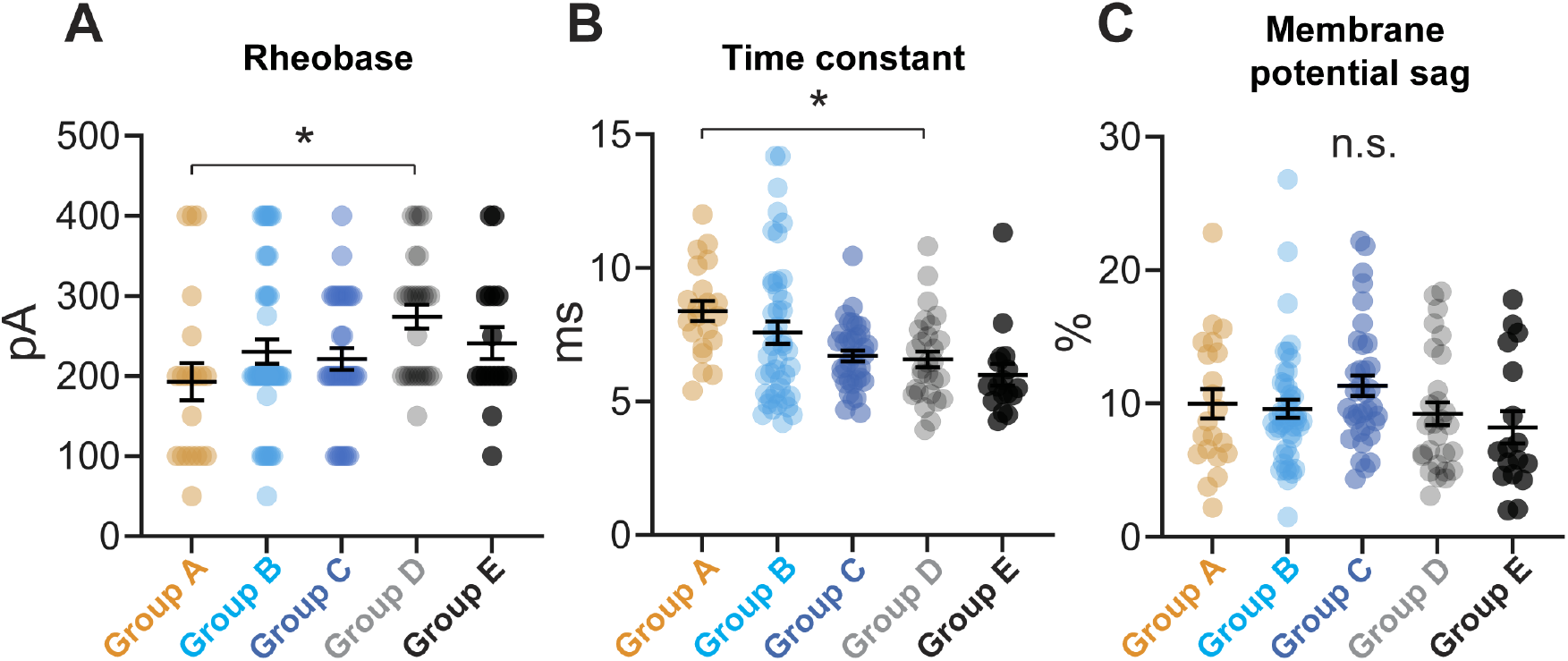
Passive membrane properties of parvalbumin interneurons. (A-C) Passive membrane properties for PV-INs from each group, including rheobase (A), membrane time constant (B), and membrane potential sag (C). See also Table S1.

**Figure S4:**
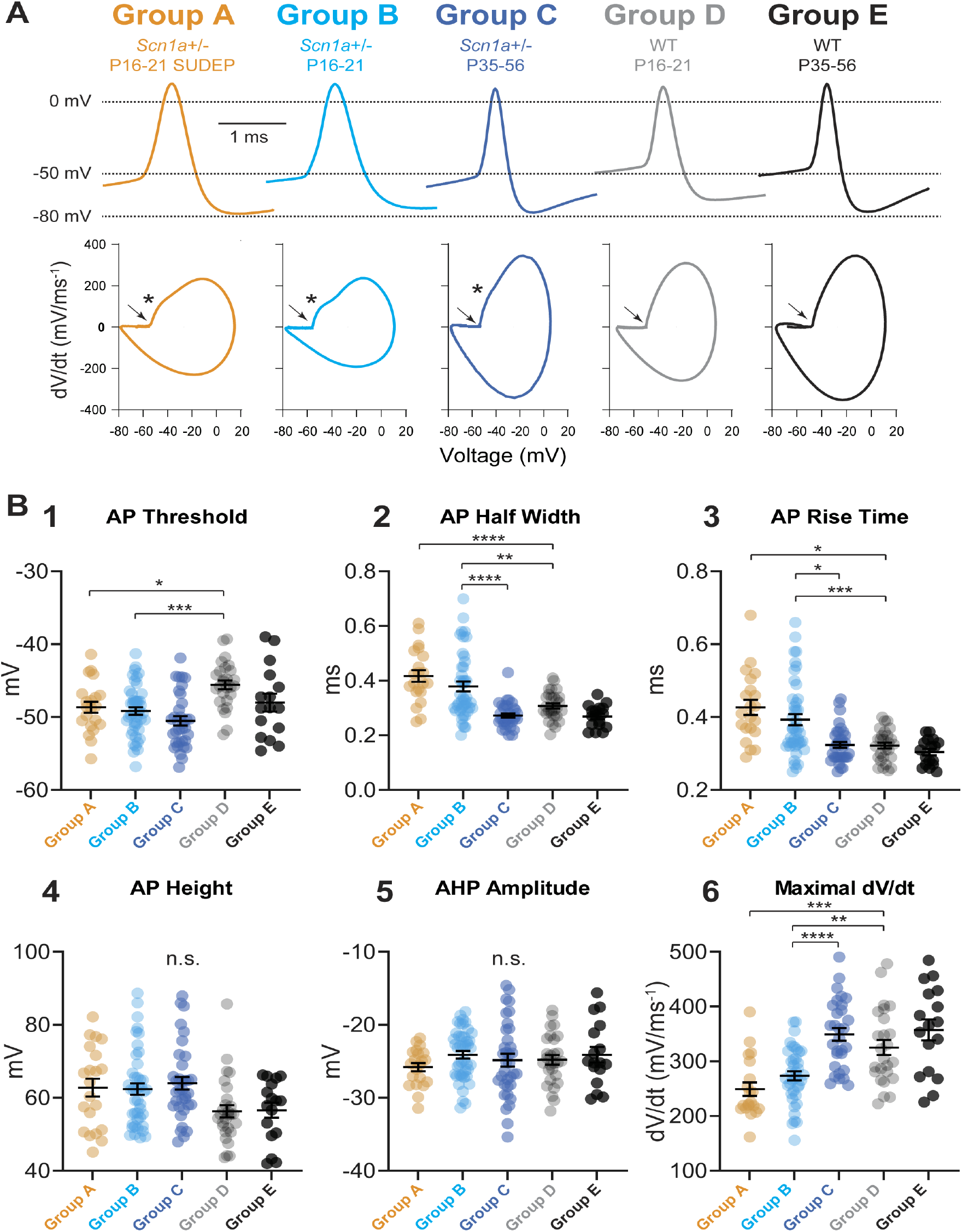
Properties of single PV interneuron action potentials. (A) Representative individual APs (top) and corresponding phase plots (bottom) from PV-INs in each group. AP threshold is indicated with an arrow; a second inflection point (“hump”), corresponding to the onset of the AIS spike, if present, is indicated by an asterisk (*). See Table S3. Note the presence of a hump in Groups A-C but not D-E. (B) Summary data for AP threshold (1), AP half-width (2), AP rise time (3), AP height (4), AHP amplitude (5), and maximal dV/dt (6). See also Table S3.

**Figure S5:**
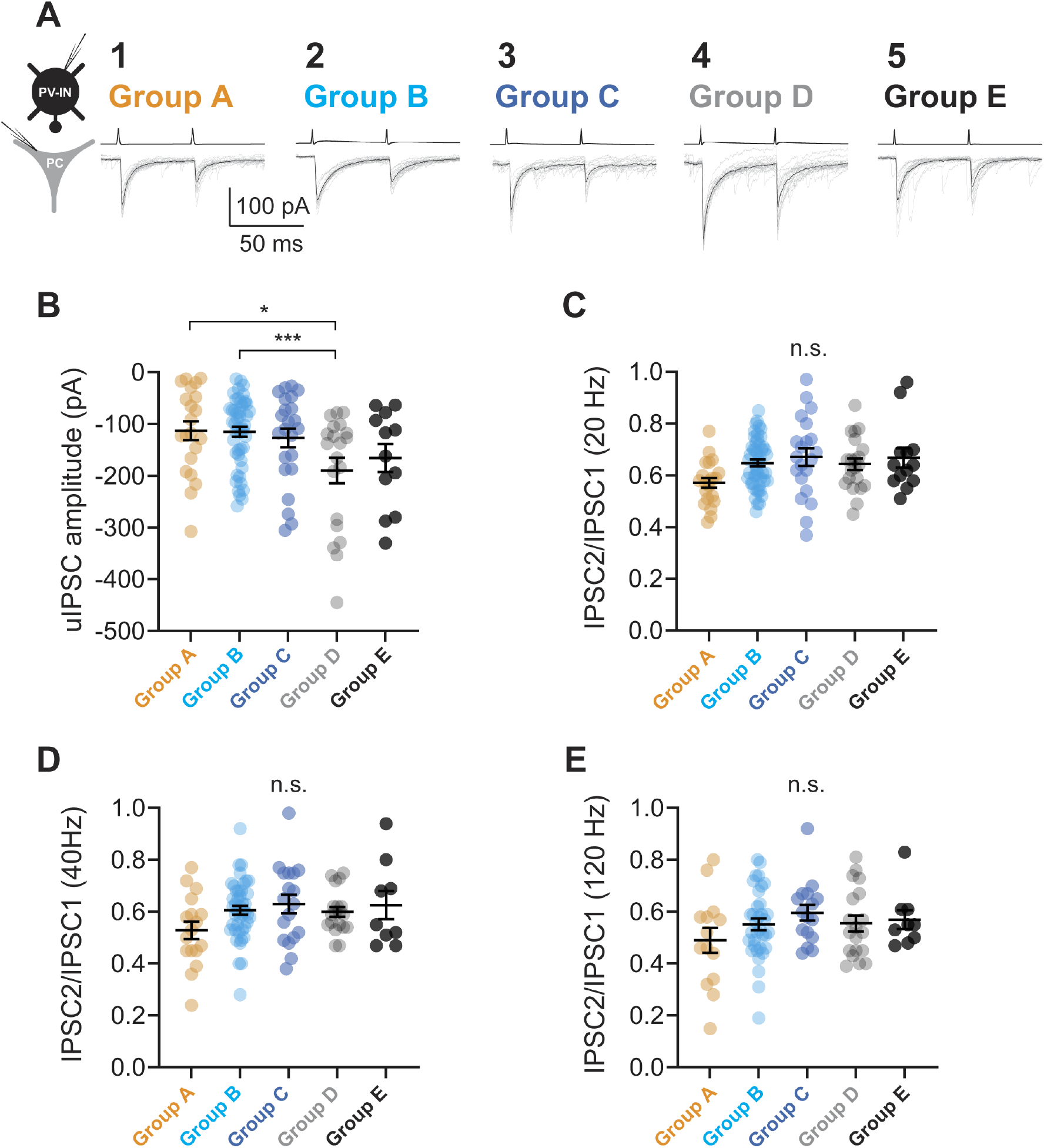
Properties of PV-IN:PC unitary connections. (A) Representative recordings of PV-IN:PC uIPSCs for Groups A-E (1-5). (B) Summary data for uIPSC amplitude. See also Table S4. (C-E) Summary data for paired-pulse ratio at 20 (C), 40 (D), and 120 Hz (E). See also Table S4.

**Figure S6:**
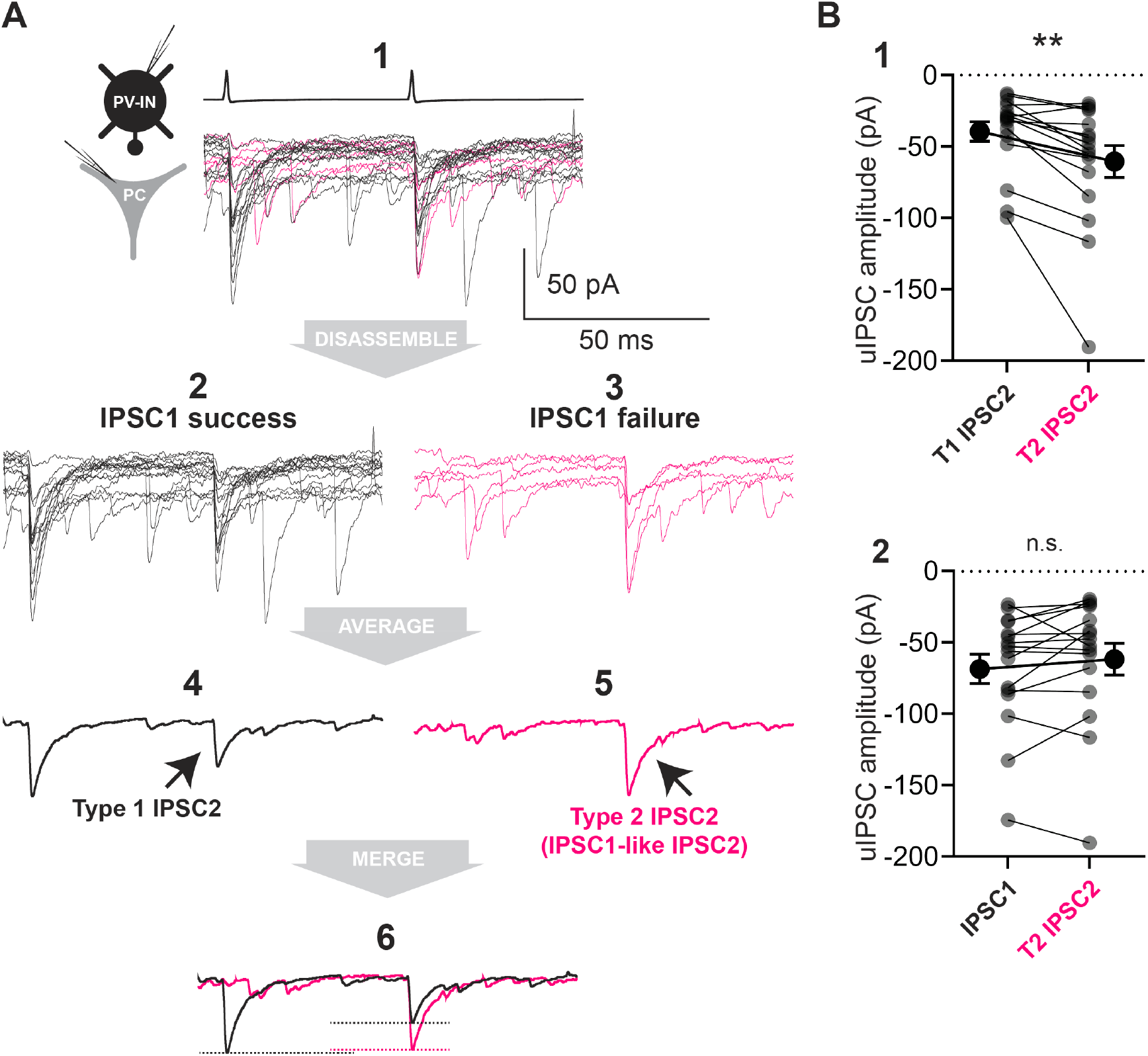
Contingency analysis of PV interneuron synaptic transmission. (A) Representative raw (light) and average (dark) traces of a PV-INs:PC unitary connection from an Scn1a+/− mouse. Shown is a set of 20 traces (A1). Black traces (15 traces; A2) indicate IPSC1 success and magenta traces (5 traces; A3) indicate failure of IPSC1. The dark black trace in A4 indicates the average of A2 while the dark magenta trace in A5 indicates the average of the traces in A3, with the merge of A4 and A5 overlaid in A6. We defined IPSC2 following success of IPSC1 as “Type 1 (T1) IPSC2,” and IPSC2 following failure of IPSC1 as “Type 2 (T2) IPSC2”. (B) Summary data for spike contingency analysis. Comparison of the mean amplitude of T1 IPSC2 and T2 IPSC2 (B1) and of the amplitude between IPSC1 and T2 IPSC2 (B2). Each data point is an individual cell/connection. Data includes Groups A, B, and C.

**Figure S7:**
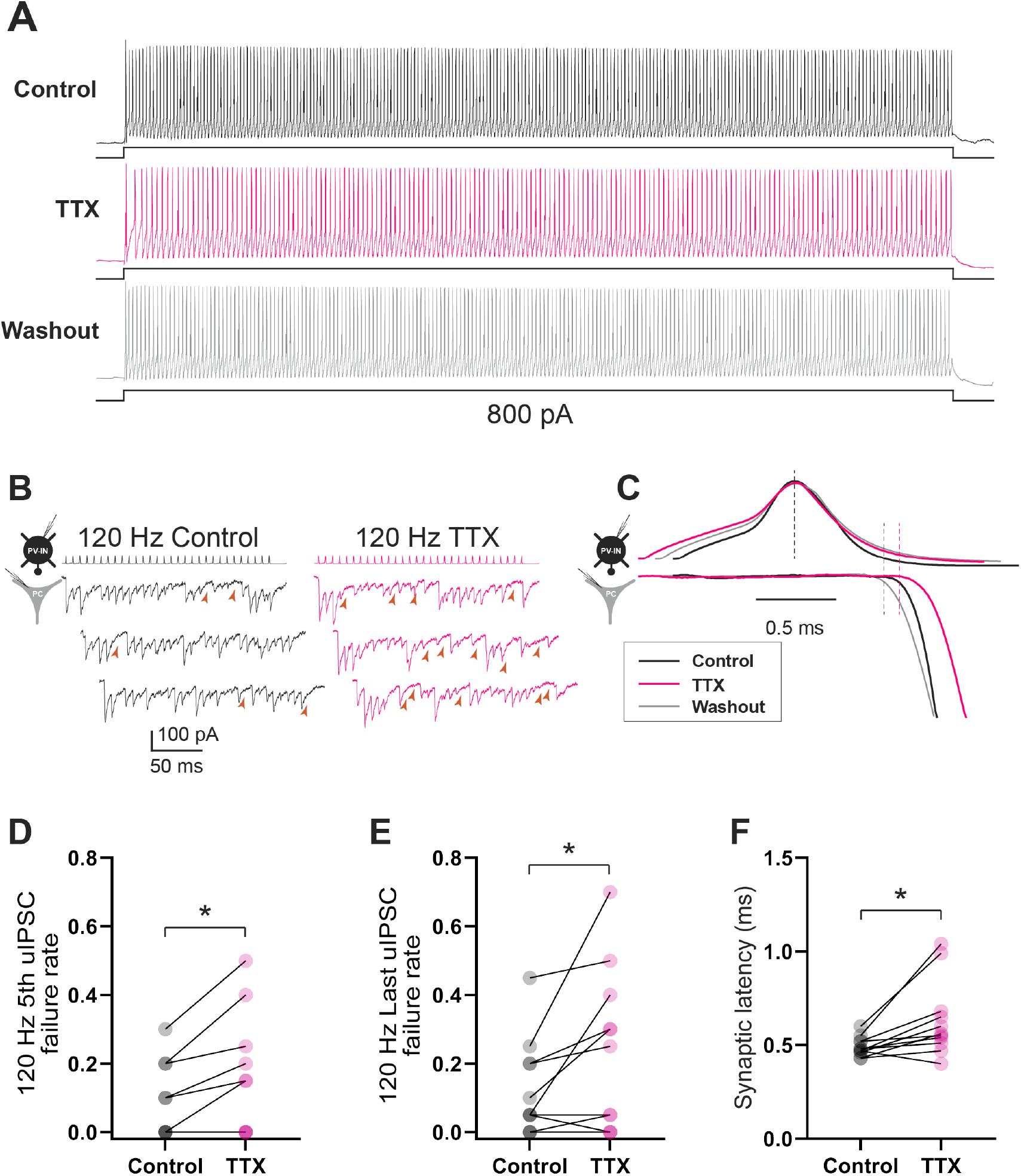
Low concentrations of TTX reproduce the effect of heterozygous loss of Nav1.1 on PV-IN synaptic transmission. (A) Electrophysiological discharge pattern of a PV-IN from a P18 WT.PV-Cre.tdT mouse in response to a 600-ms 800 pA current injection before (top, black) and after (middle; magenta) bath application of 10 nM TTX, with washout (bottom; gray). (B) Repetitive stimulation of presynaptic PV-INs elicited IPSCs in postsynaptic principle cells (left) with a higher failure rate after bath application of 10 nM TTX (right). Note that the presynaptic PV-IN can follow 120 Hz stimulation in the presence of TTX. (C) Prolonged synaptic latency after application of TTX. Shown is a presynaptic action potential aligned to the peak (top) with the onset of the postsynaptic IPSC response (bottom). (D-F) Summary data showing increased failure rate for the 5th (D) and last (E) IPSC in a train at 120 Hz and for synaptic latency (E).

**Figure S8:**
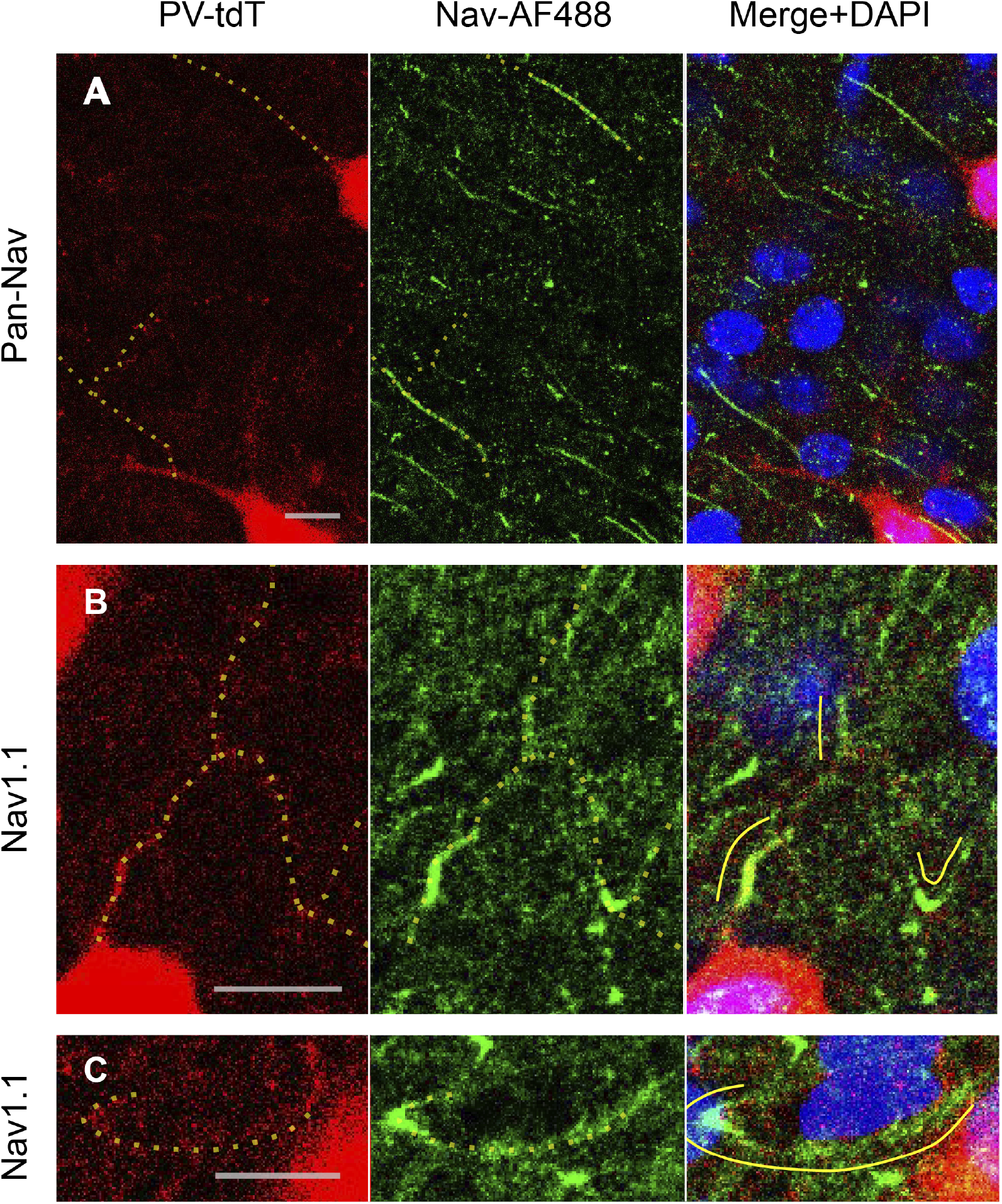
Immunohistochemical analysis demonstrates Nav1.1 and Pan-Nav labelling of parvalbumin interneuron axons. (A) Example confocal image taken from an adult WT.PV-Cre.tdT mouse, demonstrating immunoreactivity for Pan-Nav (green) as well as endogenous tdTomato signal (red) in layer IV barrel cortex. Pan-Nav labelling shows some smaller puncta, but mainly labels longer linear structures, some of which colocalize to PV-IN AIS (yellow dotted lines). (B-C) Immunohistochemistry for Nav1.1 performed as in Figure S8A. Note that Nav1.1 expression at PV-IN AIS is also apparent, but labels a shorter, more proximal extent of axon relative to pan-Nav, as reported previously. In addition, in layer IV, there is dense neuropil labelling, which partially colocalizes with PV-IN branch points (B) and presumptive basket cell pre-terminal axon/terminal forks (C). Scale bar is 5 *µ* M for all images.

**Figure S9:**
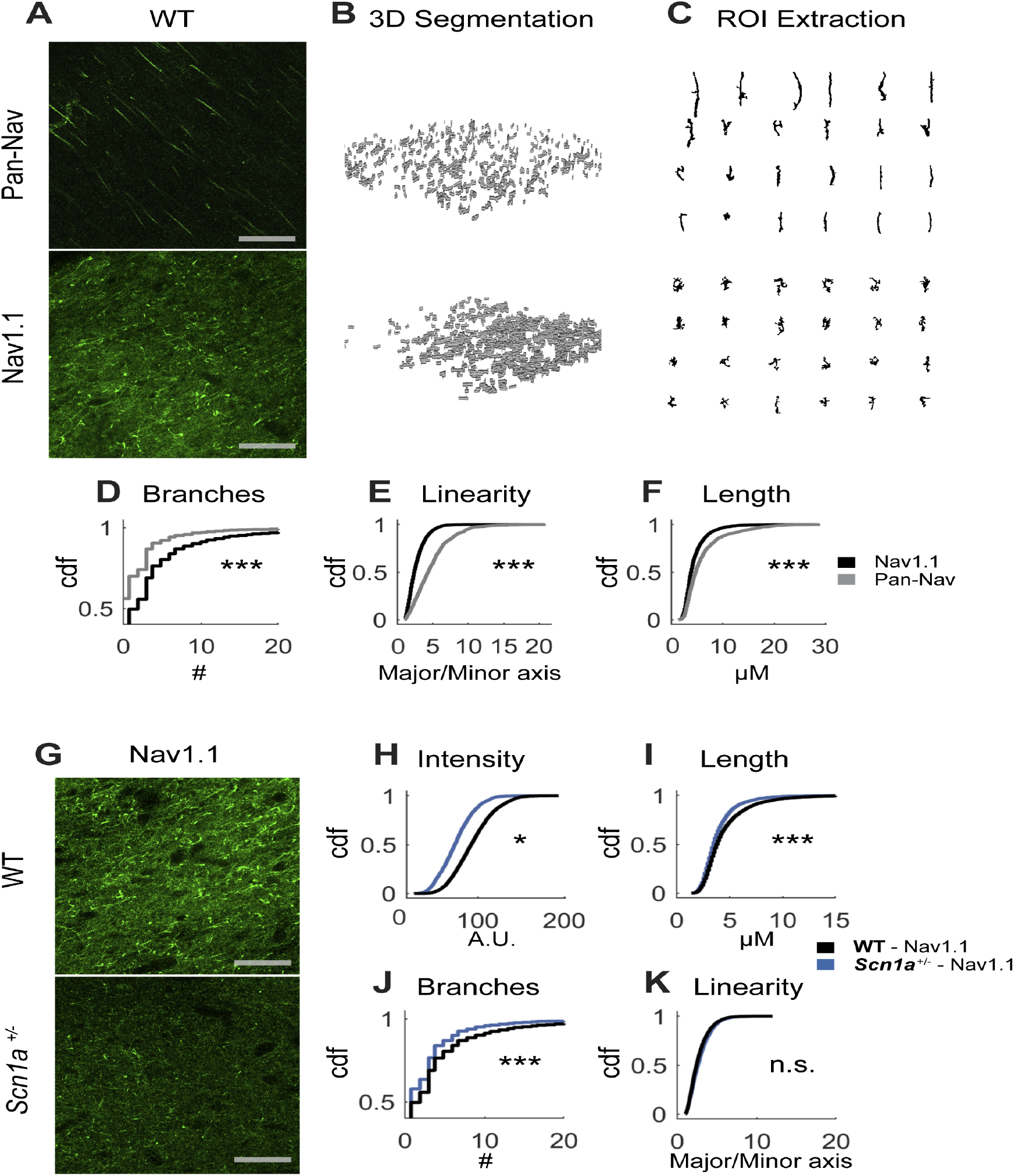
Comparison of Pan-Nav and Nav1.1 labelling in WT and Scn1a+/− mice. (A) Quantification of both Pan-Nav and Nav1.1 labelling (scale bar = 40 *µ* M) using (B) automated filtering and segmentation of reconstructed z-stacks of layer IV barrel cortex to extract (C) individual Nav puncta. Nav1.1 is more likely to label complex and/or branching structures compared to Pan-Nav (D-F). Quantification of the number of branch points, linearity, and major axis length of Nav1.1 and Pan-Nav puncta (6718 puncta from 10 Nav1.1 z-stacks, and 1409 puncta from 7 Pan-Nav z-stacks from 3 WT mice). (G) Example confocal images of Nav1.1 labelling in Layer IV barrel cortex from both WT and Scn1a+/− mice (H-K). Using the same segmentation as above, quantification of the normalized intensity of extracted puncta, as well as the parameters in D-F (6718 puncta from 10 z-stacks from 3 WT mice, and 3130 puncta from 10 z-stacks 3 Scn1a+/− mice; *, p<0.05,**, p<0.01, ***, p<0.001 whether there is a difference in means between groups determined by a mixed effects linear model treating mice as random effects).

**Figure S10:**
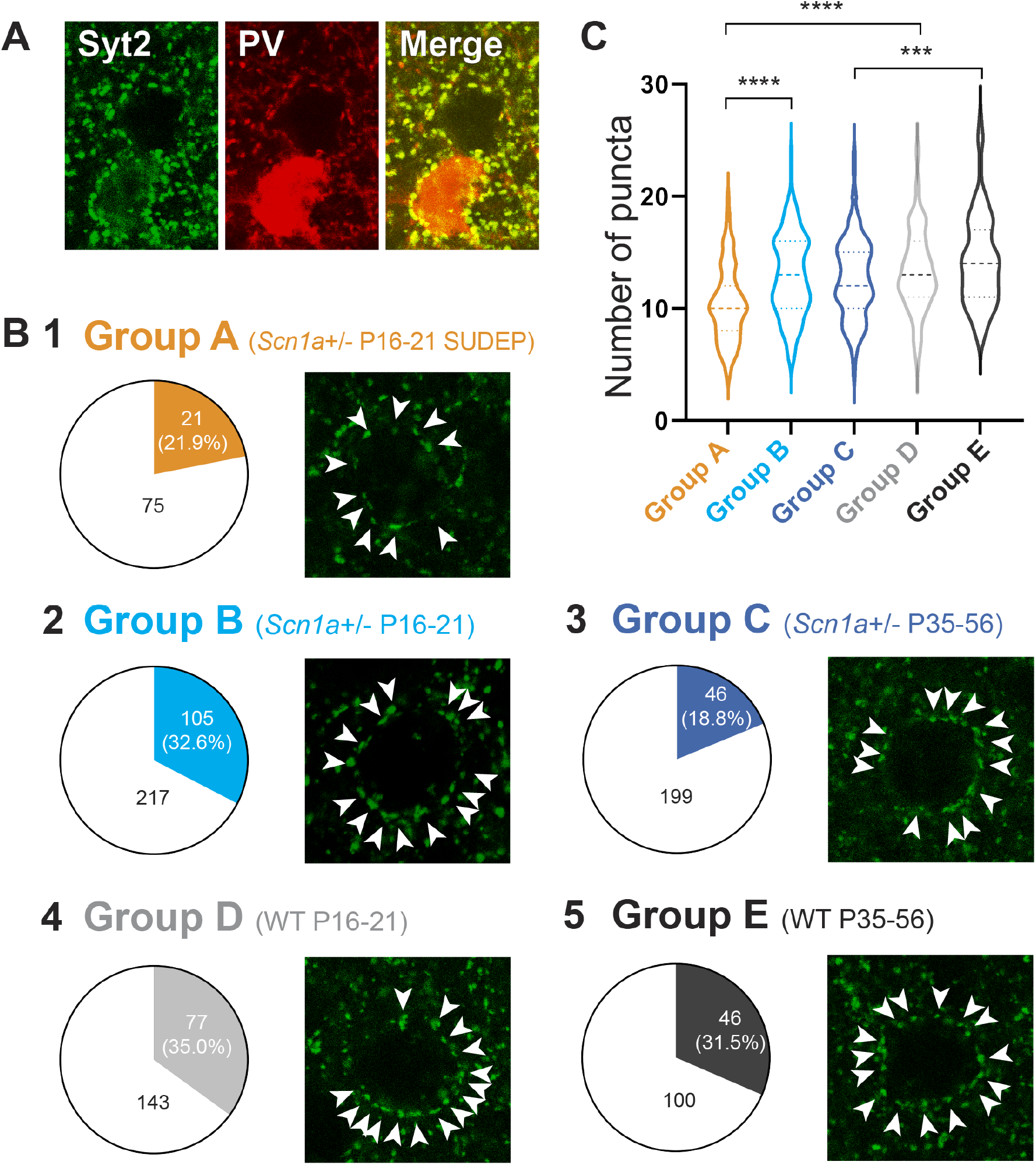
Lower PV interneuron connection probability and decreased bouton numbers in Scn1a+/− mice. (A) Shown are confocal images from an Scn1a+/− mouse at age P55 (i.e., Group C) indicating immunoreactivity for Syt2 (left; green) and tdTomato expression in Scn1a.PV-IN.tdTomato mice (middle; red), with merge (right). (B) Representative confocal images of perisomatic Syt2-positive PV-IN synaptic boutons (arrowheads) surrounding principle cells in each Group A-E, with pie charts depicting the PV-IN:PC connection probability for Groups A-E in multiple whole-cell recording experiments in acute brain slice. (C) Violin plots illustrating summary data for the number of perisomatic puncta per principle cell across Groups A-E.

**Figure S11:**
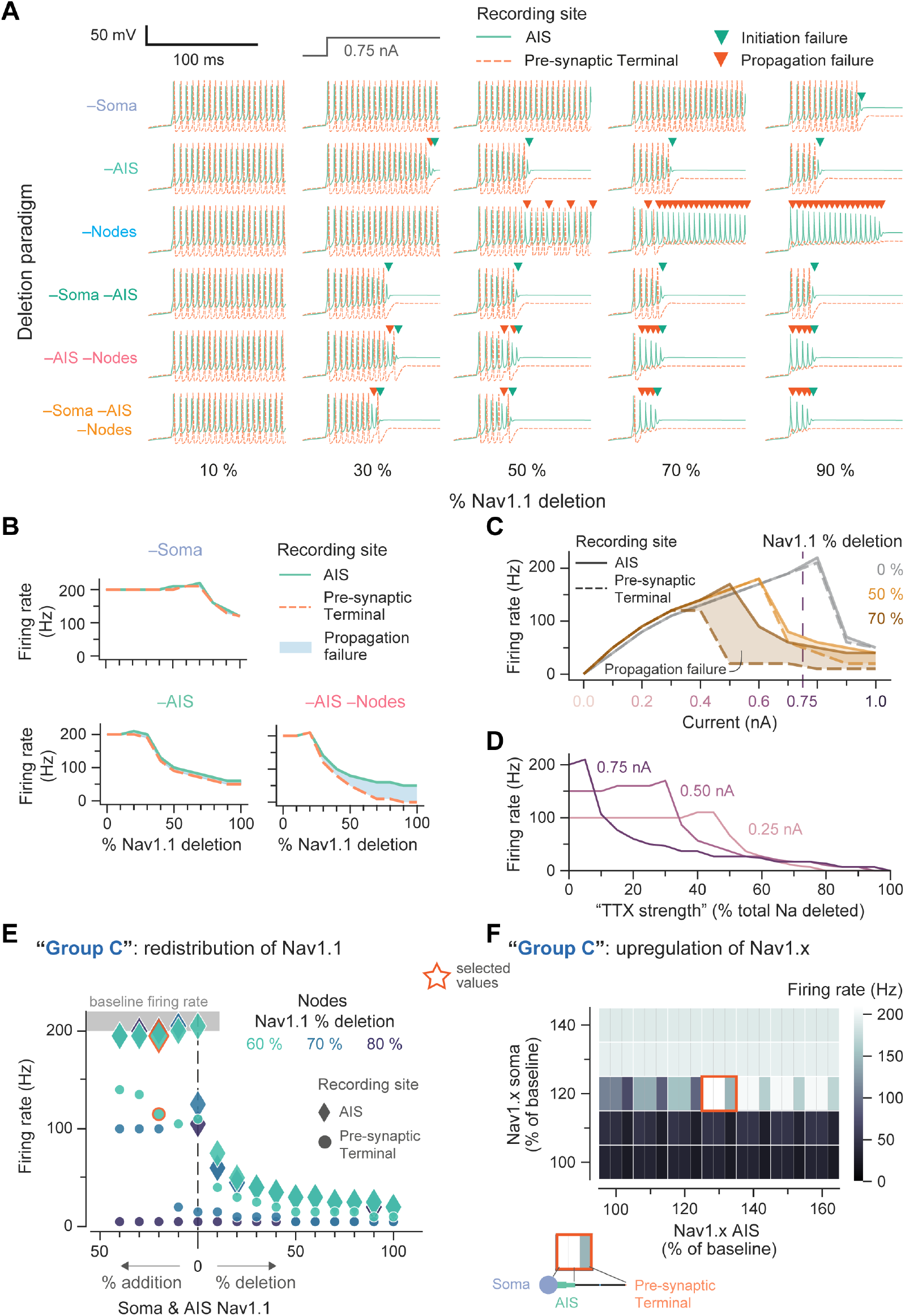
Model parameters used to explore the mechanism of the observed experimental results. (A) Membrane potential traces recorded at the axon initial segment (AIS, solid light green line) and pre-synaptic terminal (dashed light orange line), following an injection of 0.75 nA current into the soma from 20 ms. Colored downward arrows represent the time of a failure (initiation, dark green, or propagation, dark orange), the expected time that an action potential would occur. Columns represent 10, 30, 50, 70, and 90 % extent of Nav1.1 deletion (proportional to a neuronal compartment’s baseline). Rows are the different deletion paradigms explored. (B) The average firing rate of a neuron with progressive Nav1.1 deletion. The difference in firing rates between the AIS (solid light green line) and the pre-synaptic terminal (dashed light orange line) represents the propagation failure (shaded blue area), which increases with further deletion. (C) Input-output curves for baseline (0 %), 50 %, and 70 % Nav1.1 deletion. The difference in firing rates between the AIS (solid light green line) and the pre-synaptic terminal (dashed light orange line) represents the propagation failure (shaded blue area). Note that the interneuron enters depolarization block at higher input currents, as seen experimentally. Propagation failure is exacerbated by Nav1.1 deletion. The dashed purple line is 0.75 nA, which is the input current typically used elsewhere. (D) For a control, all the sodium channels in the model were modulated, rather than just the Nav1.1 channel, as a representation of TTX strength. The interneuron firing rates for different input currents were evaluated against total sodium deletion to assess their drop-off. (E) A possible mechanism that Group C interneurons used to recovery their action potential generation to baseline levels was to redistribute Nav1.1 channels from the nodes to the AIS and soma. The firing rate at the AIS (diamond markers) and pre-synaptic terminals (circle markers) was explored to investigated as a function of Nav1.1 redistribution to elicit similar baseline firing rate (gray shaded area) at the AIS but still exhibiting impaired propagation. To reach this behavior, Nav1.1 conductances in the soma and AIS were above baseline levels to compensate for the (further) nodal Nav1.1 deletion (60 %, light green; 70 %, blue; and 80 %, dark blue). Note that the total Nav1.1 conductance was still below baseline levels; that is, the total additional Nav1.1 conductance in the soma and AIS was less than Nav1.1 deleted at the nodes, for the range explored. The values chosen for the traces in Figure 6 are indicated by orange outlines: 20 % addition of soma and AIS Nav1.1, and 60 % deletion of nodal Nav1.1. (F) Another possible mechanism that Group C interneurons used to recover their action potential generation to baseline levels was to upregulate other sodium channels like Nav1.3 or Nav1.6 while Nav1.1 was 50 % deleted. Here, the model’s other transient sodium channel “Nav1.X”, as previously described in (Berecki et al. 2019), was upregulated to recover activity. Each block (white border) in the heatmap shows the firing rate for the soma, AIS, and pre-synaptic terminals from left to right (separated by light gray lines). To recover activity, the Nav1.X could be 120 % of baseline at the soma and 130 % of baseline at the AIS, as represented by the orange border and used in Figure 6.

**Figure S12:**
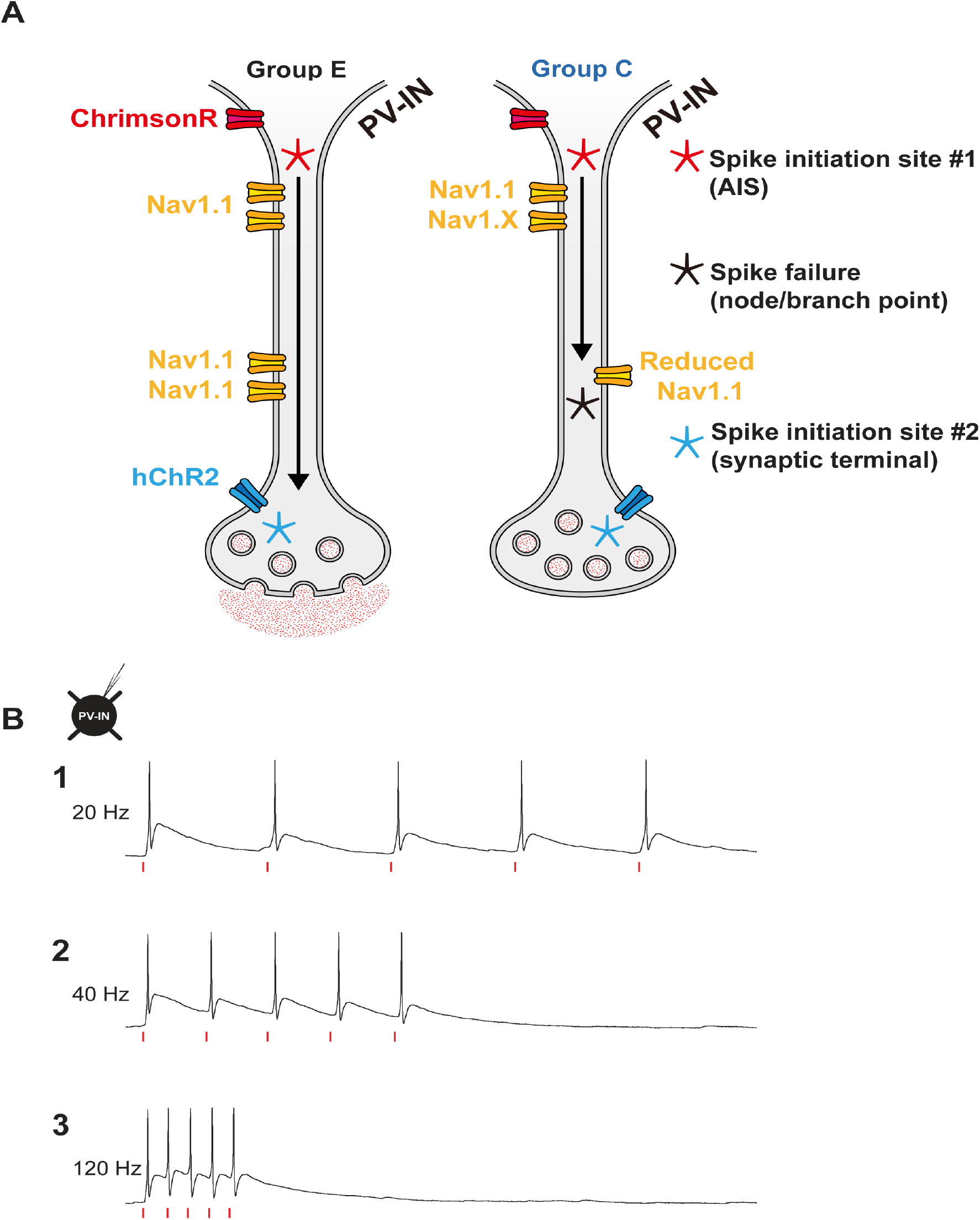
Analysis of PV interneurons across development. (A) Five experimental groups were studied: Scn1a+/− mice recorded at P16-21 and deceased by P35 (Group A; orange); Scn1a+/− mice recorded at P16-21 that survive to P35-56 (Group B; light blue); Scn1a+/− mice recorded at P35-56 (Group C; dark blue); wild-type littermates at P16-21 (Group D; gray) and P35-56 (Group E; black). (B) The mini-slice procedure. Mini-slices (B1-3) were generated from biopsies of primary somatosensory neocortex of pre-weanling (P16-21) Scn1a.PV-Cre.tdTomato and age-matched WT.PV-Cre.tdT mice. Acute brain slices (B4) were generated using standard techniques at later time points (P35-56) from Scn1a+/− mice that did not die prior to P35 (Group C) and from age-matched wild type littermates at P35-56 (Group E).

**Figure S13:**
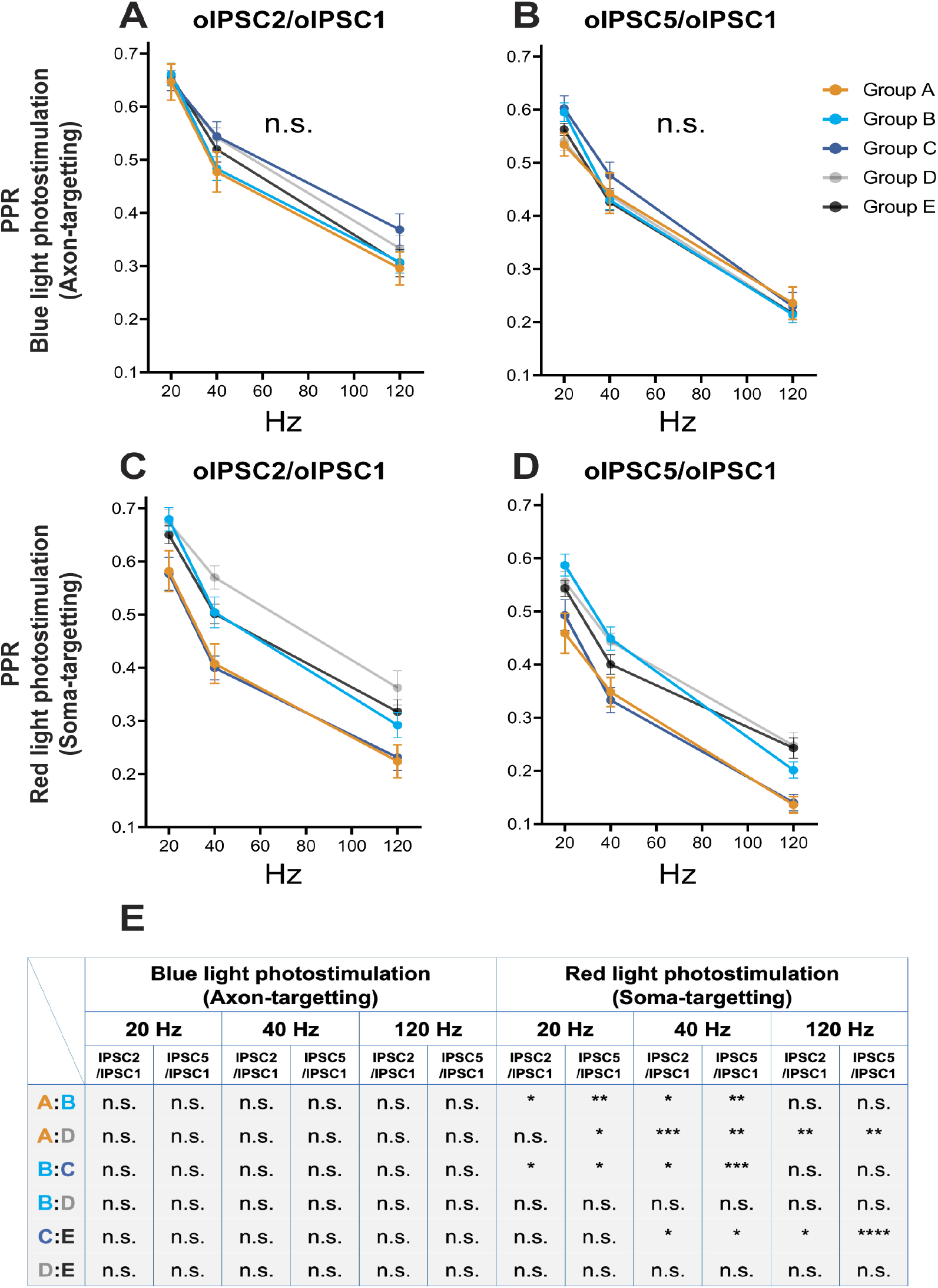
Photoactivation of hChR2 and ChrimsonR reliably drive action potentials in PV interneurons across a range of frequencies. (A) Schematic illustrating a PV-IN expressing 2 types of excitatory opsin at spatially distinct sites. (B) A whole-cell current-clamp recording from a PV-IN with representative traces for red light-driven trains of five action potentials at 20 (1), 40 (2), and 120 Hz (3).

**Table S1:**
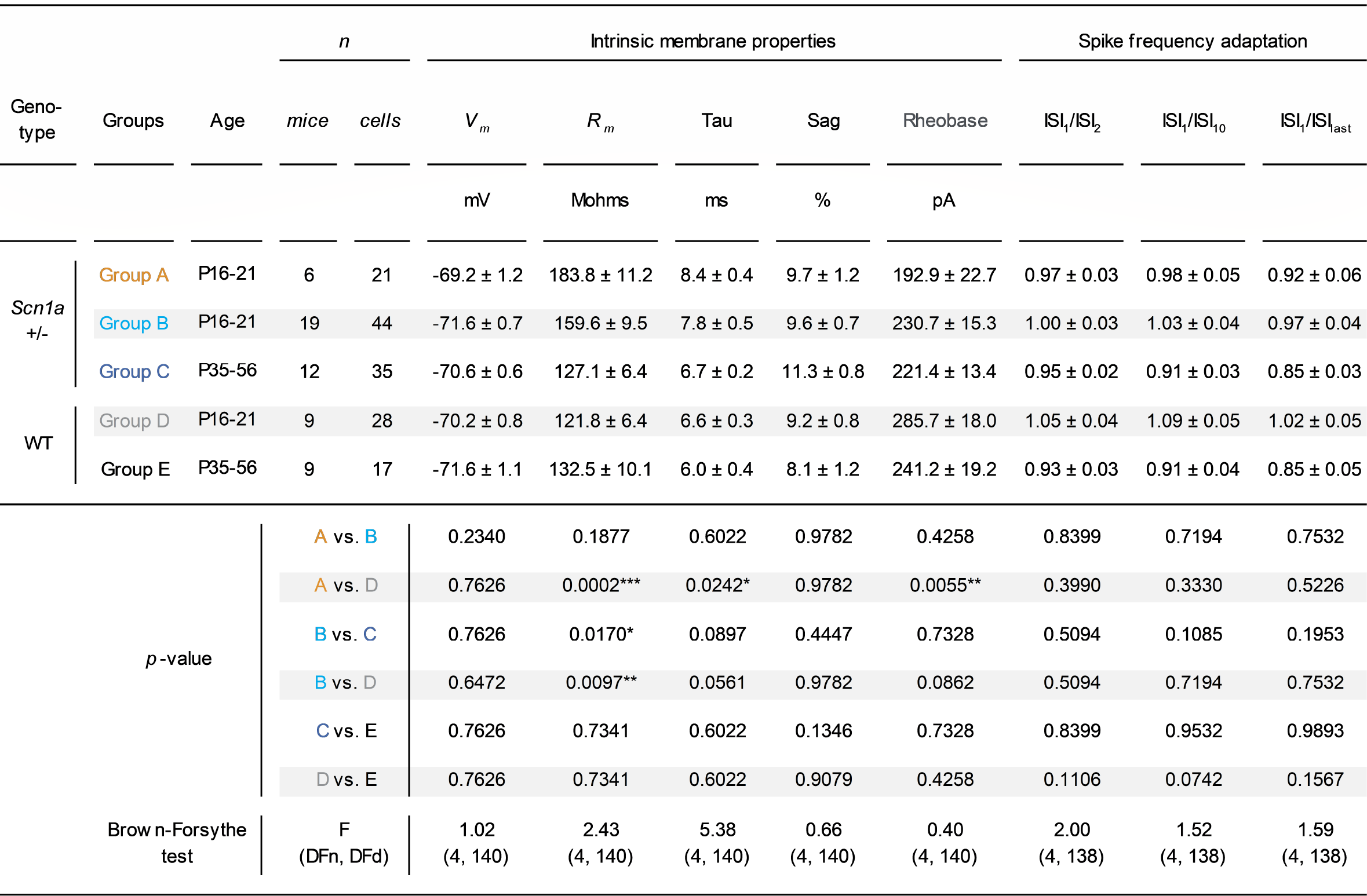
Electrophysiological properties of neocortical PV interneurons. p < 0.05; ** p < 0.01; *** p < 0.001; via one-way ANOVA with post-hoc Holm-Sidak Test between genotype or age.

**Table S2:**
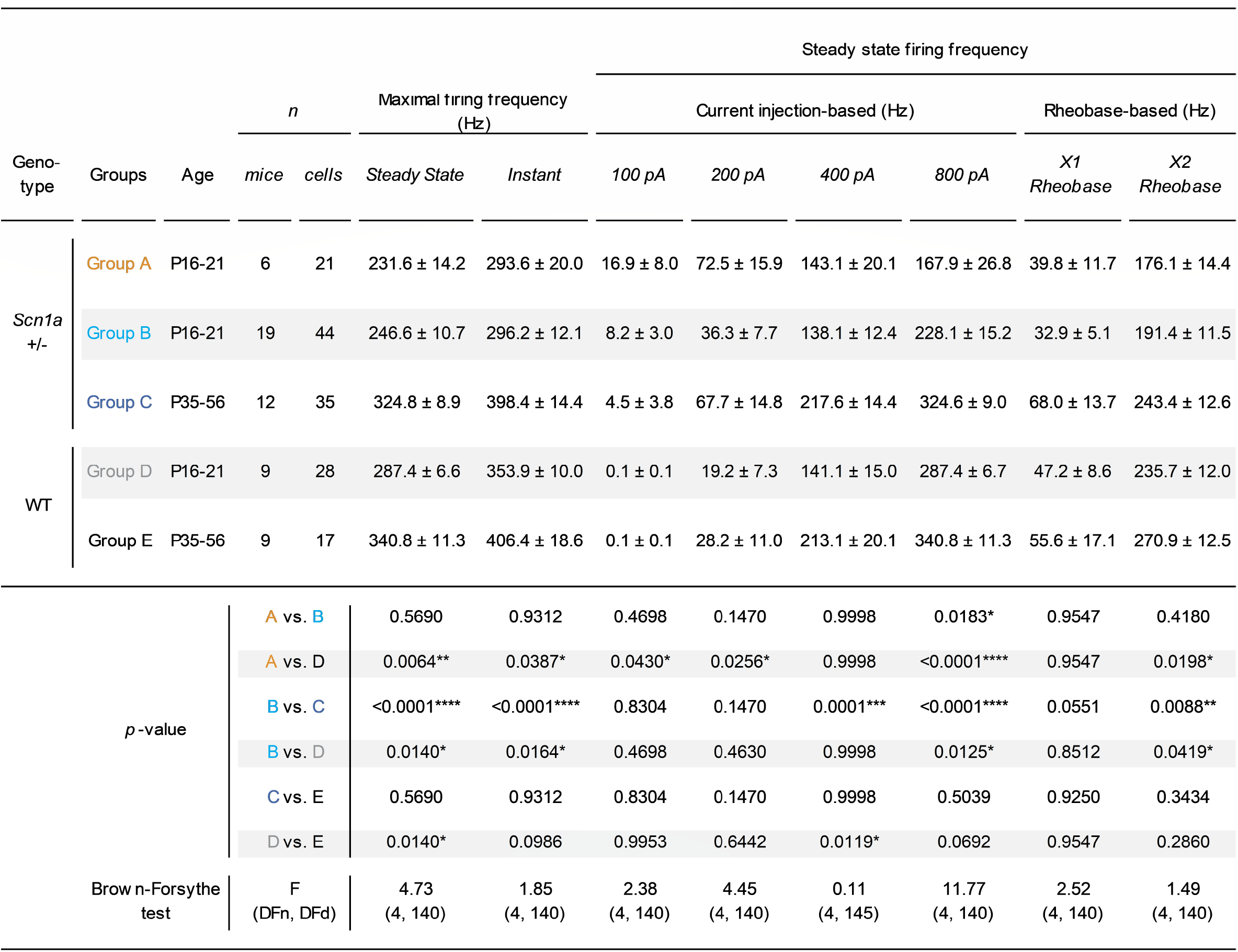
Firing frequency of neocortical PV-INs in each group. p < 0.05; ** p < 0.01; *** p < 0.001; **** p < 0.0001; via one-way ANOVA with post-hoc Holm-Sidak Test between genotype or age.

**Table S3:**
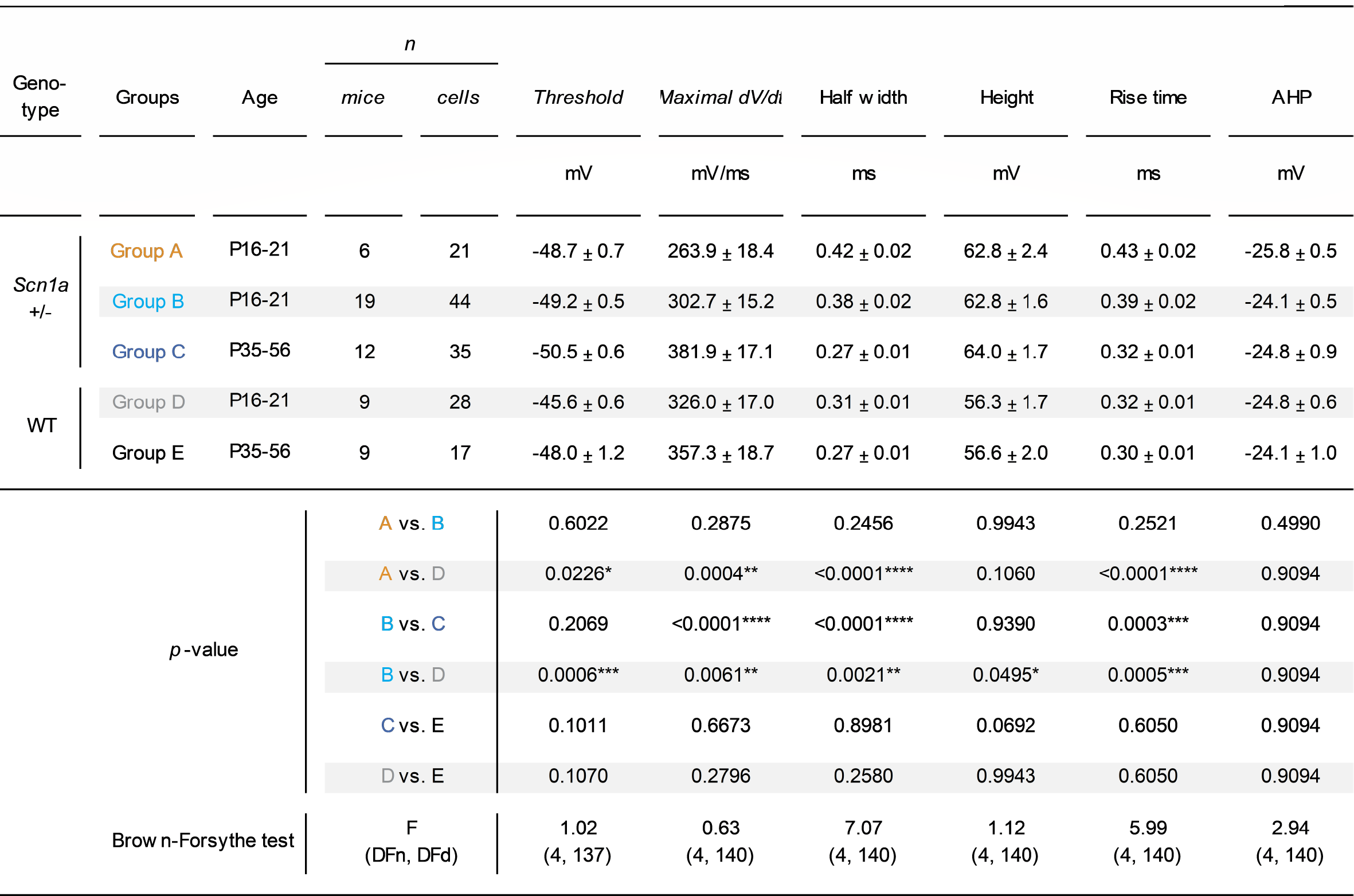
Electrophysiological properties of single action potentials in neocortical PV interneurons. p < 0.05; ** p < 0.01; *** p < 0.001; **** p < 0.0001; via one-way ANOVA with post-hoc Holm-Sidak Test between genotype or age.

**Table S4:**
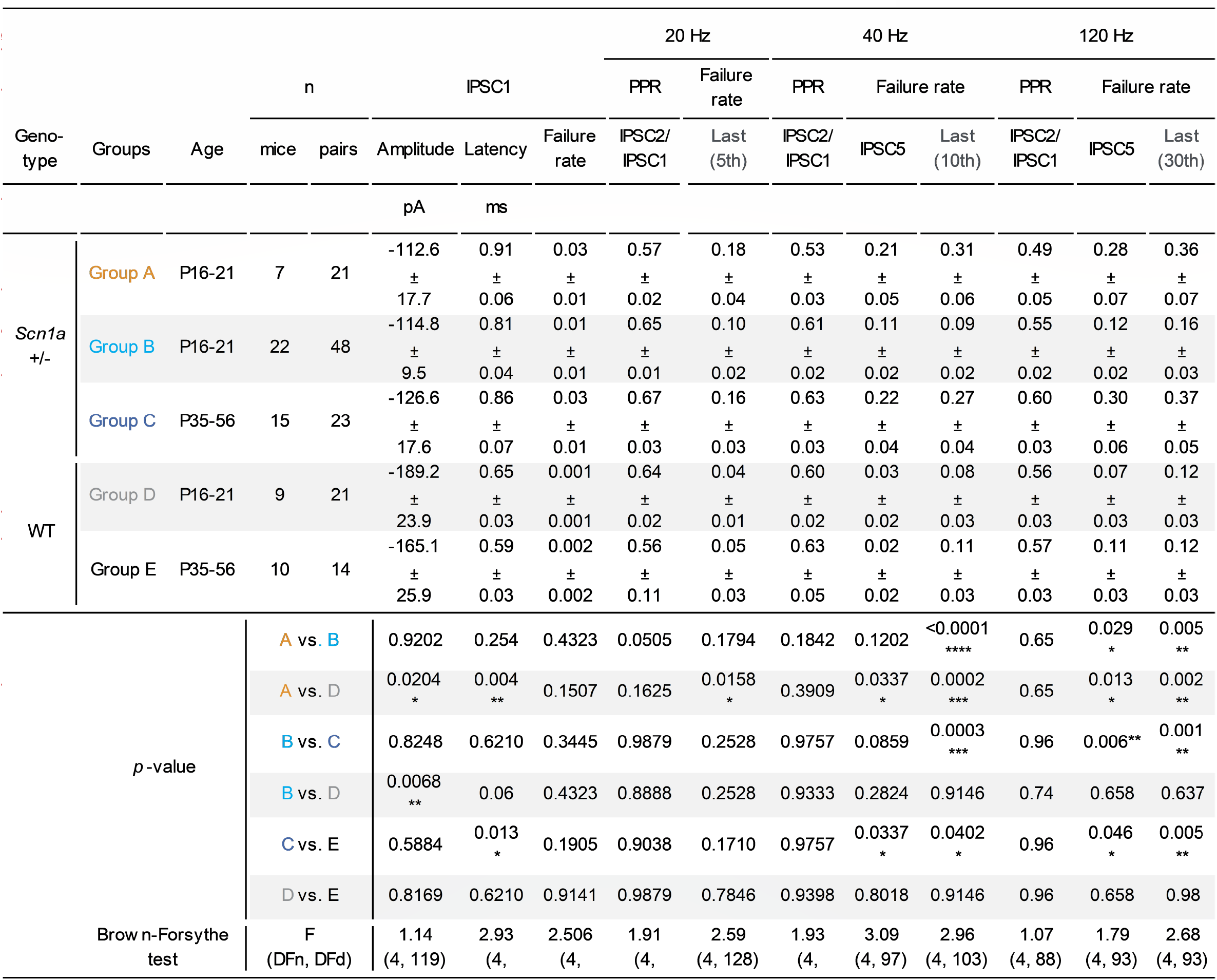
Unitary synaptic properties of PV interneuron:principal cell connections. p < 0.05; **p < 0.01; *** p < 0.001; **** p < 0.0001; via one-way ANOVA with post-hoc Holm-Sidak Test between genotype or age.

**Table S5:**
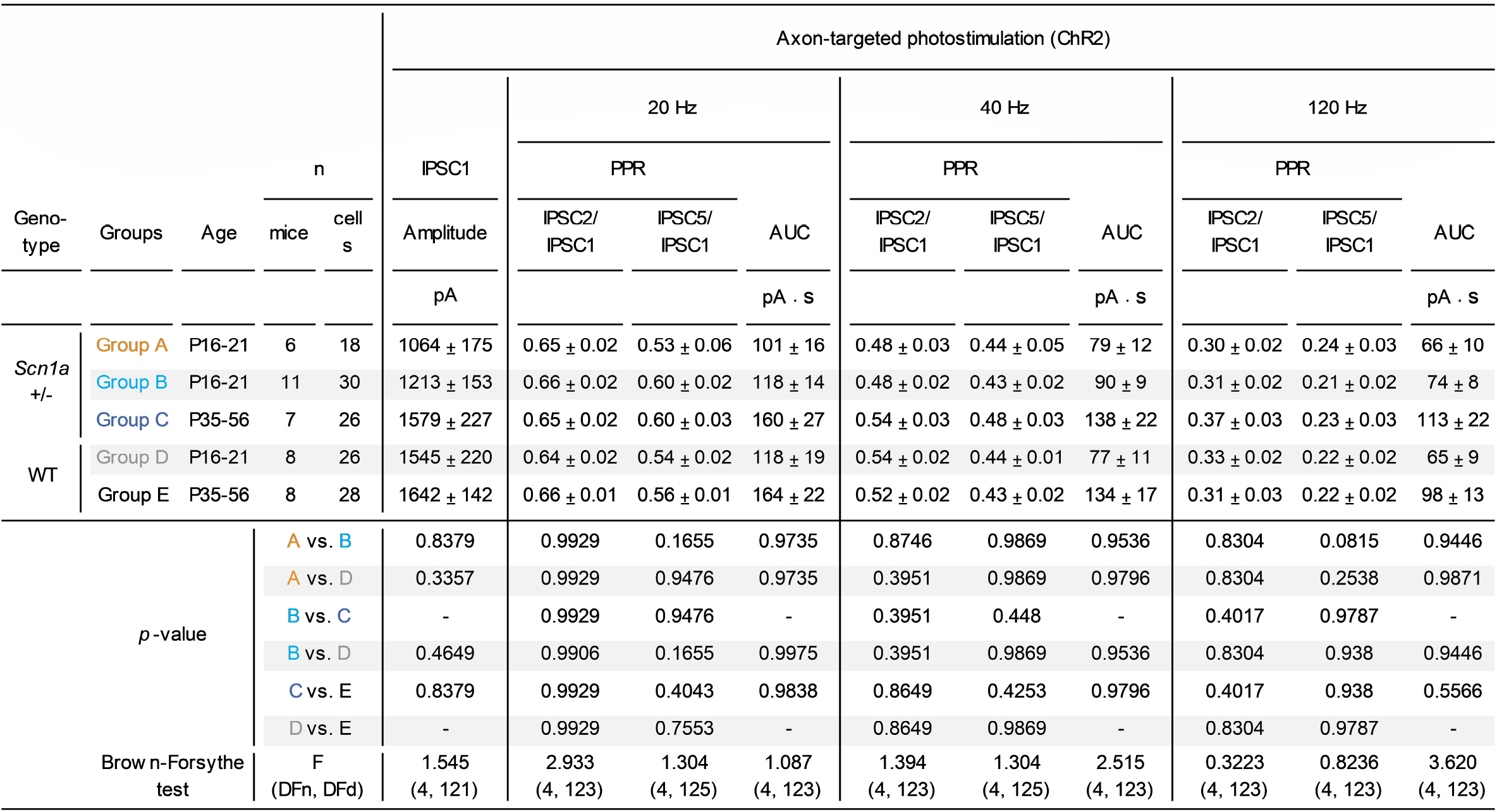
Properties of ChR2-evoked synaptic transmission. Shown is oIPSC data recorded in principle cells in response to blue light photostimulation of ChR2 expressed in surrounding PV interneurons. Analysis was via one-way ANOVA with post-hoc Holm-Sidak Test between genotype or age.

**Table S6:**
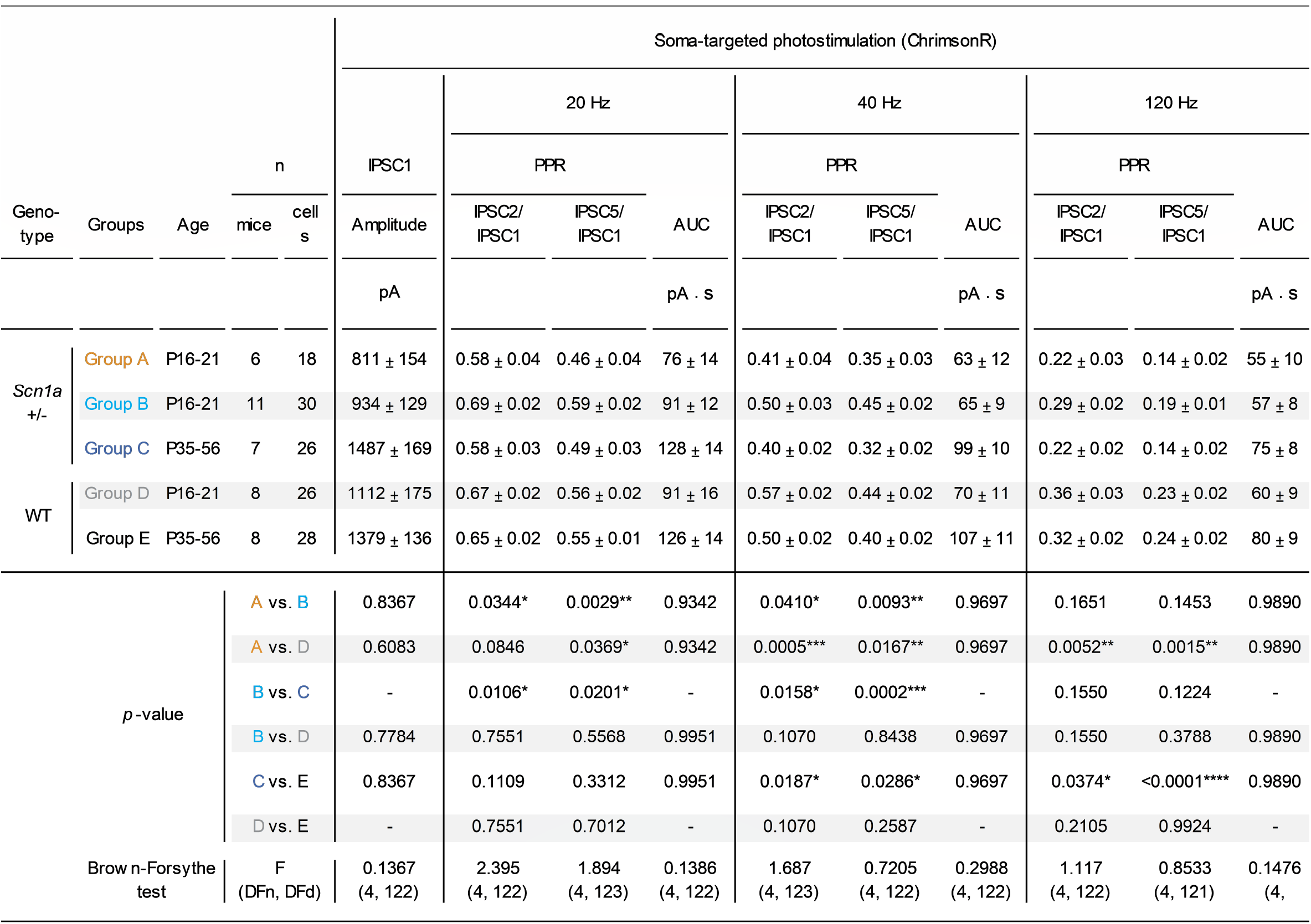
Properties of ChrimsonR-evoked synaptic transmission. Shown is oIPSC data recorded in principle cells in response to red light photostimulation of soma-targeted ChrimsonR.mRuby2.ST expressed in surrounding PV interneurons. * p < 0.05; ** p < 0.01; *** p < 0.001; **** p < 0.0001; via one-way ANOVA with post-hoc Holm-Sidak Test between genotype or age. Red (660 nm) photostimulations were recorded from PCs.

**Table S7:**
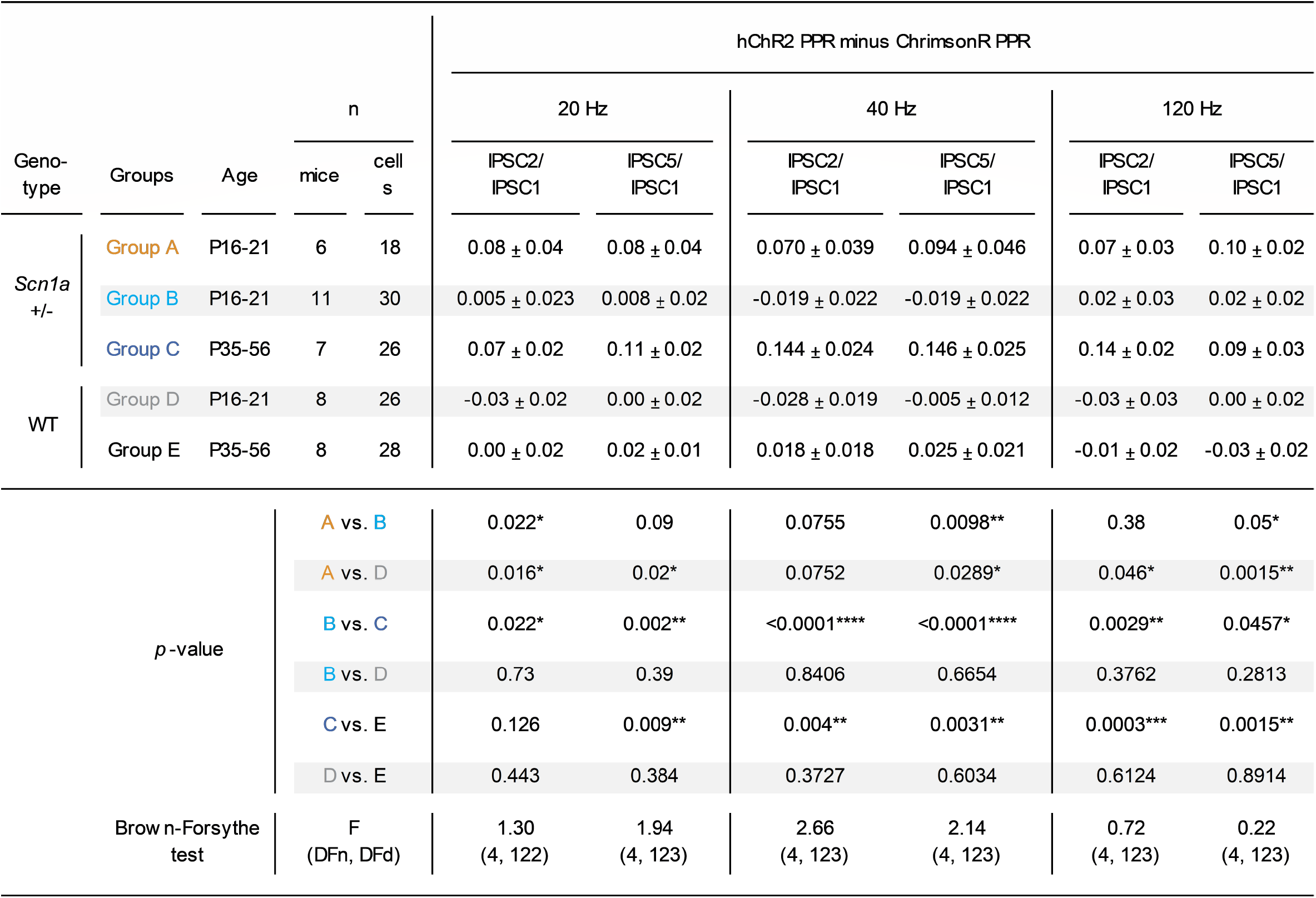
hChR2 (axon-expressed) PPR minus ChrimsonR (soma-targeted) PPR. p < 0.05; ** p < 0.01; *** p < 0.001; **** p < 0.0001; via one-way ANOVA with post-hoc Holm-Sidak Test between genotype or age. Both photostimulations were recorded from the same cells.

